# Single Amino Acid Disease Mutations Act as Genetic Glues to Drive Neo-substrate Degradation

**DOI:** 10.64898/2026.09.17.752235

**Authors:** Baiyun Wang, Chi-Wei Yeh, Hsuan-Mei Lin, Huigang Shi, Haibin Mao, Shu-Chuan Chen, Kun-Hai Yeh, Chen-Hsin Yu, Ning Zheng, Hsueh-Chi S. Yen

## Abstract

Disease-associated missense mutations are well known to affect protein turnover by either impairing substrate-ubiquitin ligase interactions or inducing misfolding-triggered protein clearance. Whether such mutations can drive gain-of-function in substrate-selective protein degradation remains under-explored. Here, we develop a tri-color global protein stability system to profile mutational effects across the intrinsically disordered regions (IDRs) of human transcription factors. Unexpectedly, we identify single amino acid substitutions that create functional short linear motif degrons to drive targeted protein degradation independently of protein quality control. These disease variants are exemplified by the gain of an FRY-like degron in GATA2, whose deficiency leads to impaired immune function. Structural analyses reveal that the substituted residue enables neomorphic interaction between the transcription factor and the KLHL15 E3 ligase by complementing their otherwise suboptimal interface. The prevalence of such glue-like mutations beyond transcription factors suggests that quasi-degrons are widespread in protein IDRs and can be functionalized by simple mutations, a finding that may inform the rational discovery of molecular glue degraders.

## INTRODUCTION

The ubiquitin-proteasome system (UPS) provides a sophisticated mechanism to regulate cellular protein abundance with high precision. In human cells, this precision is achieved through hundreds of ubiquitin E3 ligases, which function at the terminal step of a three-enzyme cascade to label specific substrate proteins with proteasome-targeting polyubiquitin chains.^1,2^ In coordination with molecular chaperones, a subset of these E3 ligases plays a role in protein quality control by recognizing and ubiquitinating misfolded proteins or orphan subunits of protein complexes.^3^ The majority of E3 ligases, however, recognize signature structural features of their cognate substrates in the form of short linear motifs (SLiMs), secondary structures, or tertiary and quaternary folds.^4,5^ These structural elements, collectively known as degrons, are often exposed or modified by cellular cues, including post-translational modifications (PTMs) or changes in the functional states of the E3 or substrate, to enable conditional and signal-responsive protein degradation. Remarkably, naturally occurring and synthetic small molecules can act as molecular glues to cement E3-degron interface and promote productive substrate ubiquitination and degradation.^6–9^ The large number of E3 ligases encoded by the human genome implies that a correspondingly diverse repertoire of degrons is continuously surveilled by the UPS to maintain protein homeostasis and coordinate responses to a wide range of cellular signals.

The scale and pervasive function of the UPS in human cells predict its central involvement in disease. To date, disease mutations are known to dysregulate the UPS through two broad mechanisms. First, they can disrupt the protein–protein interactions that underlie E3–substrate recognition, thereby stabilizing proteins that would otherwise be degraded. Such dysregulation is best illustrated by cancer-associated missense mutations in the NFE2L2 (NRF2) degrons and the phospho-degron binding site of the E3 ligase FBXW7.^10,11^ Second, disease mutations can compromise protein folds and expose them to UPS-based quality control, accelerating their degradation. For instance, substitution of a single amino acid in the DNA-binding domain of p53 can trigger its misfolding and subsequent degradation, reducing the abundance of this critical tumor suppressor.^12^

Beyond these well-established mechanisms, pathogenic genetic variations can also co-opt substrate-selective E3 ligase activity to promote gain-of-function protein degradation. Overexpression of ubiquitin ligases such as MDM2, SPOP, and KLHL24, for example, has been shown to promote enhanced or aberrant substrate turnover in cancer and genetic skin disorders.^13–15^ More strikingly, insertion-deletion mutations in the E3 ligase KBTBD4 found in medulloblastoma resurface its substrate-binding domain to engage members of the class I histone deacetylases, HDAC1 and HDAC2, thereby capturing their interacting partners, the transcriptional co-repressor CoREST and the histone demethylase LSD1, as neo-substrates for degradation.^16–18^ However, given the structural complexity of this interaction, how broadly disease mutations can enable such neomorphic E3–substrate interactions remains an open question.

Here, by systematically surveying disease-associated missense mutations in the intrinsically disordered regions (IDRs) across all human transcription factors, we unexpectedly discover that single amino acid substitutions can act as genetic glues to create functional SLiM degrons, driving targeted degradation through substrate-selective E3 ligases independently of protein quality control. The widespread occurrence of IDR-located latent degron sequences reveals an inherent susceptibility of the human proteome to mutation-induced stability perturbation via neo-degron formation, a vulnerability that also opens opportunities for genetic glue-guided discovery of molecular glue degraders.

## RESULTS

### Mapping proteostatic effects of internal peptides by iGPS

To accurately determine the impact of IDR-residing missense mutations on protein stability, we first set to develop a reporter system tailored to internal peptides harboring potential SLiM degron activity. Large-scale degron discovery efforts to date have typically relied on appending peptide libraries to the C terminus of a reporter protein (*e.g.*, GFP) and then measuring changes in reporter abundance.^19–22^ However, this strategy has important limitations. Because peptide libraries are derived from full-length proteins without considering structural accessibility, many of the identified sequences likely correspond to buried hydrophobic regions that are only recognized by the protein quality control system upon being artificially exposed to the solvent. Moreover, C-terminal fusion to GFP fails to recapitulate the native internal context of these sequences. As a result, many candidate “degrons” cannot be validated in the properly folded full-length proteins, obscuring their functional relevance.

To address these limitations, we developed iGPS (internal Global Protein Stability), a tri-color reporter system designed to better mimic the native context of internal peptides. In this system, a single expression cassette produces three fluorescent proteins (BFP, GFP and RFP) from a single mRNA transcript. RFP serves as an internal control, whereas BFP and GFP form a dual-fluorescent fusion protein connected by Gly–Ser linkers, into which the peptide of interest is inserted (BFP–GS–peptide–GS–GFP) (**Fig. 1A**). The BFP/RFP and GFP/RFP ratios provide orthogonal readouts of degron activity, minimizing false positives arising from unintended proteolytic cleavage between BFP and GFP. We confirmed that the three fluorescent signals operate independently and detected no measurable fluorescence resonance energy transfer (FRET) in the BFP–GFP fusion construct (**Fig. S1A**).

**Figure 1.**
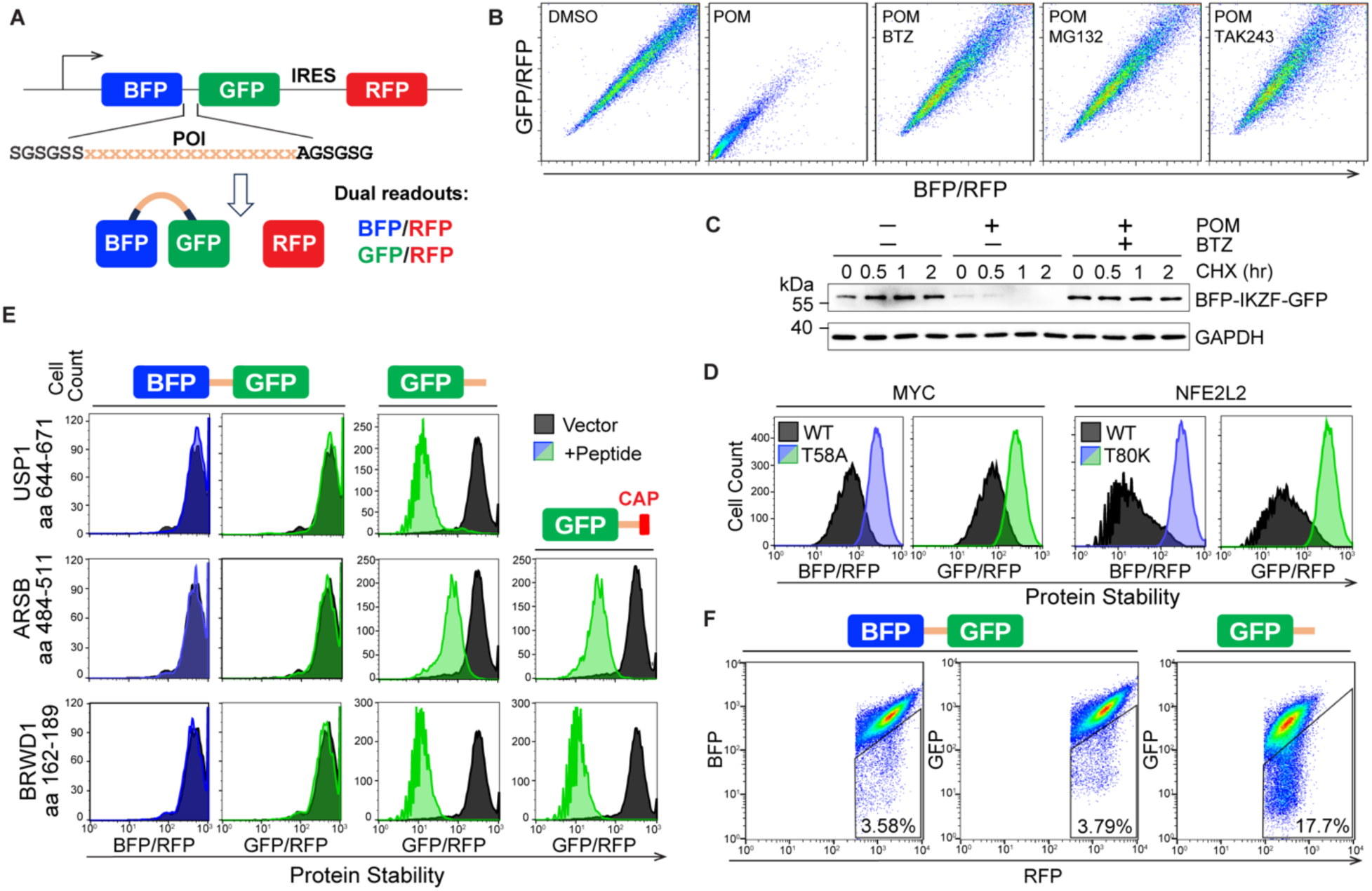
Development of iGPS to investigate the effects of disease-associated missense mutations on protein stability. (A) Schematic of the iGPS reporter. The peptide of interest (POI) is inserted between BFP and GFP via Gly–Ser linkers. A single promoter drives expression of an mRNA encoding RFP and the BFP–POI–GFP fusion protein, separated by an internal ribosome entry site (IRES), enabling proportional translation. BFP/RFP and GFP/RFP ratios serve as dual readouts of fusion protein stability. (B) Flow cytometry (FACS) analysis of iGPS reporter cells expressing the IKZF degron following treatment with pomalidomide (POM), proteasome inhibitors (bortezomib (BTZ) or MG132), or the E1 inhibitor TAK243. (C) Cycloheximide (CHX)-chase analysis of the BFP–IKZF degron–GFP fusion in the presence or absence of POM and BTZ. GAPDH serves as a loading control. (D) Assessment of degron activity in wild-type (WT) and mutant (Mut) peptides derived from MYC (aa 45–72) and NFE2L2 (aa 71–98) using the iGPS assay in HEK293T cells. (E) Evaluation of degron activity for the indicated peptides when positioned internally between BFP and GFP or fused to the GFP C terminus, with or without a 20-residue cap. (F) FACS analysis of cells expressing peptide libraries in either configuration, with each cell harboring a single construct. Percentages indicate peptides that promote degradation.

To validate iGPS, we incorporated a 25-residue immunomodulatory imide drug (IMiD)-inducible IKZF degron, along with wild-type–mutant peptide pairs derived from pathogenic variants in c-MYC and NFE2L2, which are known to impair degron function. As expected, in cells expressing the IKZF degron, both the BFP/RFP and GFP/RFP ratios decreased concordantly upon pomalidomide treatment, an effect that was blocked by proteasome or E1 inhibition (**Fig. 1B**). Cycloheximide-chase assays confirmed that this decrease in fluorescence reflects enhanced protein degradation (**Fig. 1C**). In addition, c-MYC T58A and NFE2L2 T80K mutant peptides were less effective at promoting degradation than their wild-type counterparts, which are constitutively active in the HEK293T cell (**Fig. 1D**).

We next examined whether peptide position influences degron activity. A diGly-containing USP1 internal peptide (aa 644–671) did not induce degradation when placed internally but did so when fused to the C-terminus of GFP, indicating that C-terminal fusion can artificially generate C-degrons (**Fig. 1E**, top panel). To test this effect systematically, we introduced the same peptide library (∼6,000 peptides) targeting internally located and intrinsically disordered regions (IDRs) of 424 cancer-related proteins either at the GFP C-terminus or between BFP and GFP and then measured their degron activity. Strikingly, C-terminal placement was more than four-fold more likely to induce degradation than internal insertion (17.7% versus 3.8%) (**Fig. 1F**). A previous internal degron screen attempted to mitigate artificial C-degron formation by adding a 20-amino-acid C-terminal cap.^19^ However, the degrons identified in that study failed to induce degradation in our iGPS-based assay, despite promoting degradation when fused to the GFP C-terminus with or without the cap (**Fig. 1E**, middle and bottom panels). Together, these results demonstrate that degron activity is highly dependent on structural context and underscore the importance of preserving native peptide positioning for accurate identification of functional internal degrons.

### Disruption of transcription factor degrons by disease-linked IDR mutations

To systematically examine how disease-associated missense mutations in IDRs could affect internal degrons and influence transcription factor stability, we compiled 2,081 IDR mutations, including 1,206 variants linked to genetic diseases across transcription factors in the ClinVar database and 875 cancer-associated mutations in 135 transcription factors annotated as oncogenes or tumor suppressors.^23,24^ IDRs were defined based on AlphaFold predictions (pLDDT < 85).^25,26^ An oligonucleotide library encoding 28-residue wild-type–mutant peptide pairs, with the variant centrally positioned, was cloned into the iGPS reporter and introduced into HEK293T cells via single-copy viral transduction. Cells were sorted into low-(2.5%) and high-stability (90%) fractions on the basis of BFP/RFP and GFP/RFP ratios, which serve as readouts of fusion protein stability. Oligonucleotides from both fractions were recovered and sequenced, and fold changes (log₂FC (low/high)) were calculated. Enrichment in the low-stability fraction (high log₂FC) indicates degron activity. Comparison of mutant and wild-type peptides using Δlog₂FC (Mut–WT) enabled identification of mutation-induced degron gain or loss (**Fig. 2A**). Three biological replicates showed strong correlation (**Figs. S1B, S1C**).

**Figure 2.**
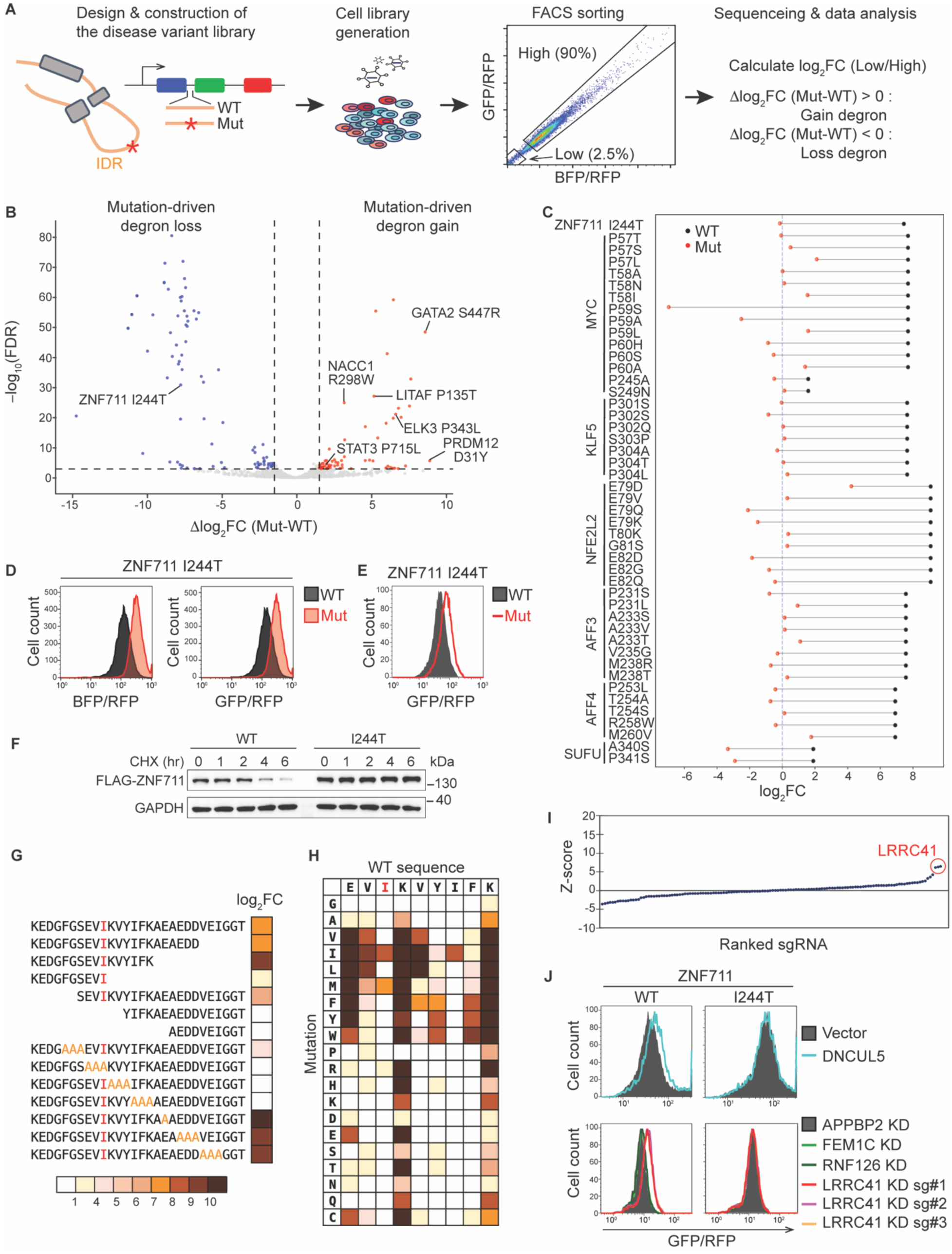
Disease-associated IDR mutations reprogram degron landscapes in human transcription factors. (A) Overview of the workflow used to identify IDR mutation-induced degron gain or loss in human transcription factors. Disease-associated variants from ClinVar and COSMIC were incorporated into peptide constructs alongside their corresponding WT sequences. The library was cloned into the iGPS reporter and introduced into HEK293T cells via single-copy lentiviral transduction. Cells were sorted into low-(2.5%) and high-stability (90%) populations on the basis of BFP/RFP and GFP/RFP ratios. Sequencing of recovered peptides enabled calculation of log₂FC (low/high) as a measure of degron activity. Comparison of WT and mutant log₂FC values (Δlog₂FC (Mut–WT)) enabled identification of mutation-induced degron gain or loss. (B) Volcano plot showing mutation-induced effects on degron activity. The x-axis represents the difference in log₂FC (low/high) between mutant and WT peptides, and the y-axis indicates statistical significance. Degron-disrupting variants are shown in blue, whereas neodegron-inducing variants are shown in red. Selected mutations are labeled. Dashed lines indicate thresholds (FDR < 0.001 and |Δlog₂FC| ≥ 1.5). (C) Dumbbell plot of representative degron-disrupting variants. (D) , (E) Validation of mutation-induced protein stabilization using peptide-based reporters (D) and full-length transcription factors (E). (F) CHX-chase analysis of FLAG-tagged full-length WT and mutant ZNF711. GAPDH serves as a loading control. (G) Mapping of the ZNF711 degron (aa 235–262) by serial truncation and alanine-scanning mutagenesis. Degron activity is shown as log₂FC, with color intensity reflecting degradation strength. (H) Saturation mutagenesis analysis of the ZNF711 core degron (aa 242–250). (I) CRISPR–Cas9 screen identifying BC-box components required for degradation of full-length WT ZNF711. Each dot represents a single sgRNA. (J) Stability of full-length WT and mutant ZNF711 in the presence or absence of dominant-negative (DN) CU5 (top panels) or following sgRNA-mediated knockout of the indicated E3 ligases (bottom panels).

Our approach identified a panel of both known and previously uncharacterized SLiM degrons. As expected, their functions can be impaired by disease-associated single missense mutations (**Fig. 2B**). Using thresholds of FDR < 0.001 and |Δlog₂FC (Mut–WT)| ≥ 1.5, we identified 79 degron-disrupting variants, including mutations in MYC, KLF5, NFE2L2, AFF3, AFF4 and SUFU (**Figs. 2C, S2A, S2B**), whose accumulation have been implicated in human diseases. These degrons are recognized by distinct E3 ligases, including FBW7 (MYC, KLF5), KEAP1 (NFE2L2), SIAH (AFF3, AFF4) and FBXL17 (SUFU),^27–33^ highlighting the robustness of the screen and the broad applicability of iGPS for identifying internal degrons.

Among the 79 degron-disrupting mutations, 37 have not been previously documented. To validate these newly identified degron-disrupting variants in a physiologically relevant context, we examined their effects in the full-length transcription factors. The germline mutation ZNF711 Ile244Thr is associated with X-linked intellectual disability, although its pathogenic mechanism remains unclear.^34^ We found that residues 235–262 in ZNF711 contains an IDR-localized degron (**Figs. 2D**, **S2C**). This missense mutation, therefore, impairs degron activity, leading to increased stability of the corresponding full-length transcription factor (**Fig. 2E, 2F**).

To further define the degradation mechanism of ZNF711, we performed serial truncation and alanine-scanning analyses, which identified a minimal nine-residue hydrophobic motif, EVI^244^KVYIFK, as the core degron (**Fig. 2G**). Saturation mutagenesis pinpointed two critical isoleucine residues (I244 and I248), with Ile244 corresponding to the disease-associated mutation site (**Fig. 2H**). A CRISPR-based screen identified the CRL5^LRRC41/MUF1^ complex as the E3 ligase targeting this degron (**Figs. 2I**). Consistently, inhibition of CRL5 ^LRRC41/MUF1^ with a dominant-negative CUL5 construct or LRRC41 knockout stabilized wild-type ZNF711 but had no effect on the Ile244Thr mutant (**Fig. 2J**).

### Disease mutation-rendered neodegrons in transcription factors

While missense mutations that disrupt degrons in transcription factors are well documented, our iGPS-based screen unexpectedly uncovered numerous gain-of-function degrons that are generated by IDR mutations. Strikingly, such neo-degron formation induced by single amino acid substitution occurred at a frequence comparable to degron loss (64 vs. 79), underscoring the prevalence of degron-like SLiMs with sub-optimal sequences in the IDRs (**Fig. 2A**). Among the variants we identified, NACC1 Arg298Trp, GATA2 Ser447Arg, PRDM12 Asp31Tyr, LITAF Pro135Thr, STAT3 Pro715Leu and ELK3 Pro343Leu stood out with variable potencies (**Figs. 3A, S3A**). The destabilizing effects of these variants were confirmed using both peptide-based assays (**Figs. 3B, S3B**) and full-length transcription factors (**Fig. 3C**). Importantly, these variants reside in IDRs and are not flanked by hydrophobic residues, suggesting that the degrons created by these variants are categorically distinct from Bag6-dependent degrons that act through protein quality control pathways.^35^

**Figure 3.**
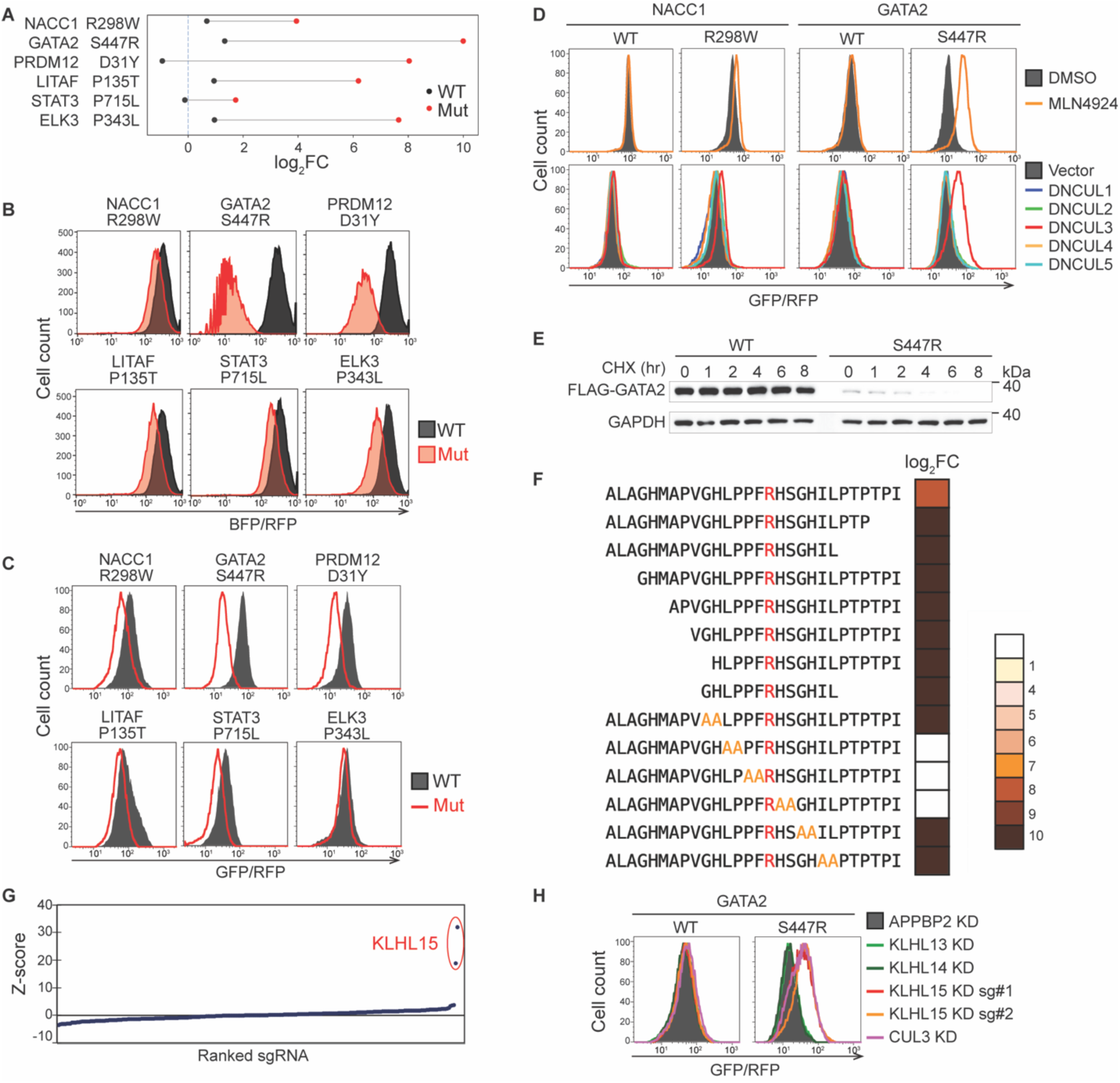
Disease-associated IDR variants induce neodegrons in human transcription factors. (A) Dumbbell plot of representative degron-generating variants. (B) ,(C) Validation of mutation-induced protein degradation using peptide-based reporters (B) and full-length transcription factors (C). (D) Protein stability analysis of full-length WT and mutant NACC1 and GATA2 in the presence or absence of the CRL inhibitor MLN4924 (top panels), or following overexpression of the indicated dominant-negative Cullins (bottom panels). (E) CHX-chase analysis of FLAG-tagged full-length WT and mutant GATA2. GAPDH serves as a loading control. (F) Mapping of the GATA2 Ser447Arg degron (aa 432–459) by serial truncation and alanine-scanning mutagenesis. Degron activity is shown as log₂FC, with color intensity reflecting degradation strength. (G) CRISPR–Cas9 screen identifying CRL3 substrate adaptors required for degradation of full-length GATA2 Ser447Arg. (H) Stability of full-length WT and mutant GATA2 following sgRNA-mediated knockout of CUL3 and associated substrate adaptors.

To further dissect this mechanism, we characterized the degradation pathways of the NACC1 Arg298Trp and GATA2 Ser447Arg mutants. Both variants acquire neodegron activity mediated by CRL3-family E3 ligases. Inhibition of CRL3 ligases by MLN4924 treatment or expression of a dominant-negative CUL3 suppresses their neodegron activities, although the destabilizing effect induced by the disease variant in GATA2 is more profound than that of NACC1 (**Fig. 3D**). Therefore, we decided to focus on the GATA2 Ser447Arg mutant.

The GATA2 Ser447Arg autosomal dominant mutation is associated with myelodysplastic syndrome and deafness–lymphedema.^36,37^ Notably, multiple independent nucleotide substitutions (c.1339A>C, c.1341C>A and c.1341C>G) converge on the same amino acid change, Ser447Arg, highlighting its clinical relevance.^36,38^ We confirmed that the Ser447Arg substitution induces a neodegron and promotes protein degradation by cycloheximide (CHX)-chase analysis using FLAG-tagged full-length GATA2 (**Fig. 3E**). Through serial truncation and alanine-scanning analyses, we identified a minimal seven-residue motif, LRRFR^447^HS, as the core degron (**Fig. 3F**), which is in the middle of the long C-terminal IDR of the transcription factor (**Fig. S3C**). A CRISPR-based screen identified KLHL15 as the substrate receptor of the CRL3 complex targeting this degron (**Fig. 3G**). Consistently, KLHL15 knockout rescues degradation of the Ser447Arg mutant but has no effect on wild-type GATA2 (**Fig. 3H**). Taken together, these findings reveal a conceptually distinct mechanism of protein loss-of-function induced by disease mutations—a single amino acid substitution within an IDR fosters a neomorphic interaction with a substrate-selective E3 ligase independently of quality control, thereby leveraging the vast UPS to actively trigger protein degradation.

### Characterization of KLHL15 and GATA2 neo-interactions

To gain deeper insight into the mechanism by which the Ser447Arg mutation (hereafter designated as S447R) drives KLHL15-depednent GATA2 degradation, we first performed co-immunoprecipitation assays in HEK293T cells to determine whether GATA2 interacts with KLHL15 *in vivo*. While wildtype GATA2 did not show association with KLHL15, its S447R mutant efficiently co-precipitated with KLHL15 and CUL3, despite expressed at a lower steady-state level (**Fig. 4A**). We further assessed the direct interaction between the GATA2 neo-degron and KLHL15 in a Ni-NTA pulldown assay using purified recombinant His-tagged KLHL15 protein and MBP-fused GATA2 peptides. Consistently, only the GATA2 S447R mutant was captured by KLHL15 (**Fig. 4B**), indicating that the disease-associated GATA2 S447R mutation enables neomorphic interactions with KLHL15.

**Figure 4.**
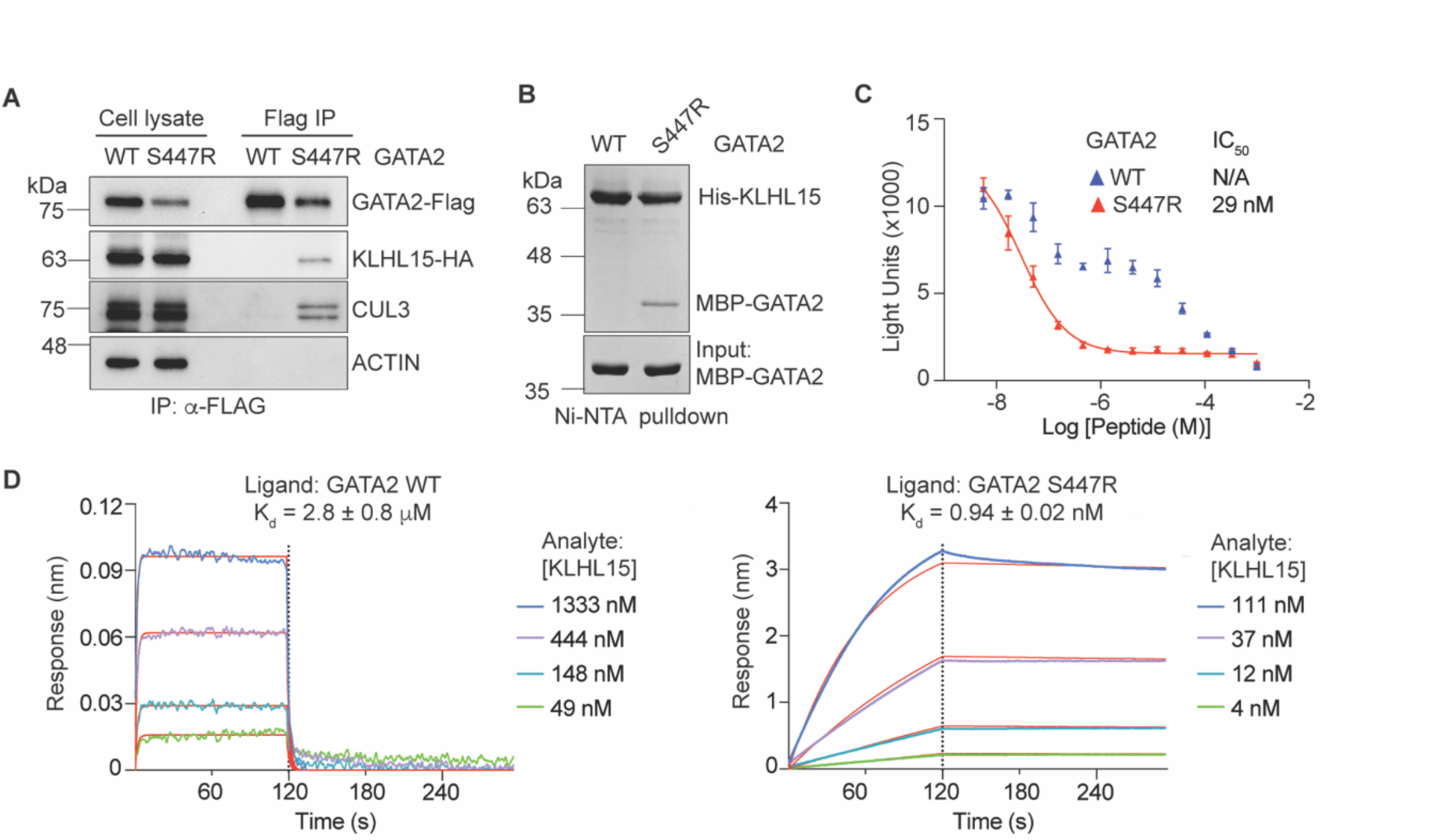
Characterization of KLHL15 and GATA2 neo-interactions. (A) Flag IP immunoblot analysis of 293T cells expressing full-length wildtype or S447R mutant GATA2-Flag. (B) Coomassie staining of *in vitro* Ni-NTA pulldown assays using recombinant His-tagged KLHL15 protein and MBP-tagged GATA2 peptides. (C) AlphaLISA competition assays for assessing GATA2 peptides binding to KLHL15. Data were presented as mean ± s.d. n = 3 biologically independent samples. IC_50_, half-maximum inhibitory concentration. N/A, not available. (D) BLI kinetic measurements of GATA2-KLHL15 interaction. Left panel, wildtype GATA2 peptide; right panel, GATA2 S447R mutant peptide. The vertical black dotted line indicates the switch from association to dissociation phase. Red lines represent global curve fitting. Kd, dissociation constant.

To quantify their binding affinity, we performed an amplified luminescent proximity homogenous assay (ALPHA)-based competition experiment. Label-free GATA2 peptides were used to compete for complex formation between affinity-tagged KLHL15 and the degron peptide of CtIP, a known KLHL15 substrate^39^ (**Fig. 4C**). In contrast to the wildtype GATA2 peptide, which poorly competed with the CtIP degron for binding KLHL15 with an atypical dose response curve, the S447R mutant peptide blocked the CtIP-KLHL15 interaction signal with an IC_50_ of 29 nM (**Fig. 4C**). Next, we carried out BioLayer Interferometry (BLI) experiments to study the binding kinetics of their interactions. Biotinylated GATA2 peptides were immobilized on streptavidin probes and recombinant KLHL15 proteins were titrated as the analyte at different concentrations. Although the BLI responses were just above the detection threshold, KLHL15 did show dose-dependent association with the wildtype GATA2 peptide. Their binding exhibited fast association and fast dissociation rates, with a calculated dissociation constant (K_D_) of 2.8 ± 0.8 μM. By contrast, the GATA2 S447R mutant peptide shows robust association with KLHL15 with an extremely slow dissociation rate. The calculated K_D_ is 0.94 ± 0.02 nM, which represents an ∼ 3,000 folds enhancement compared to the wildtype GATA2 peptide (**Fig. 4D**). Collectively, these results demonstrate that GATA2 has an intrinsic basal affinity toward KLHL15, which is biophysically detectable but physiologically insignificant. The serine-to-arginine single amino acid substitution in GATA2, remarkably, is sufficient to confer its high-affinity binding to the E3, leading to GATA2 clearance.

### Overall structure of the KLHL15-GATA2 neo-degron complex

KLHL15 has been previously characterized as a CRL3 substrate receptor that recognizes a substrate SLiM degron with a central “FRY”-like consensus motif, including FRY, FRF, LRF, and LRY.^19,39–41^ Interestingly, the neo-degron created by the GATA2 S447R mutation possess a histidine residue at the third position. Saturation mutagenesis analysis supports that FRH is indeed a variant of the “FRY” degron motif (**Fig. S3D**). To elucidate how the GATA2 S447R mutant is recognized by KLHL15, we determined the cryo-electron microscopy (cryo-EM) structure of KLHL15 in complex with the GATA2 neo-degron peptide at a resolution of 3.5 Å (**Fig. 5A**). The density of the full-length KLH15 protein is well resolved in the three-dimensional (3D) reconstitution map, enabling us to build almost the entire polypeptide with high confidence. The central eight amino acids (residue 442 to 449) of the GATA2 neo-degron also showed clear densities at the top of the KLHL15 pocket, while the flanking residues were invisible from the map, presumably in a disordered structure.

**Figure 5.**
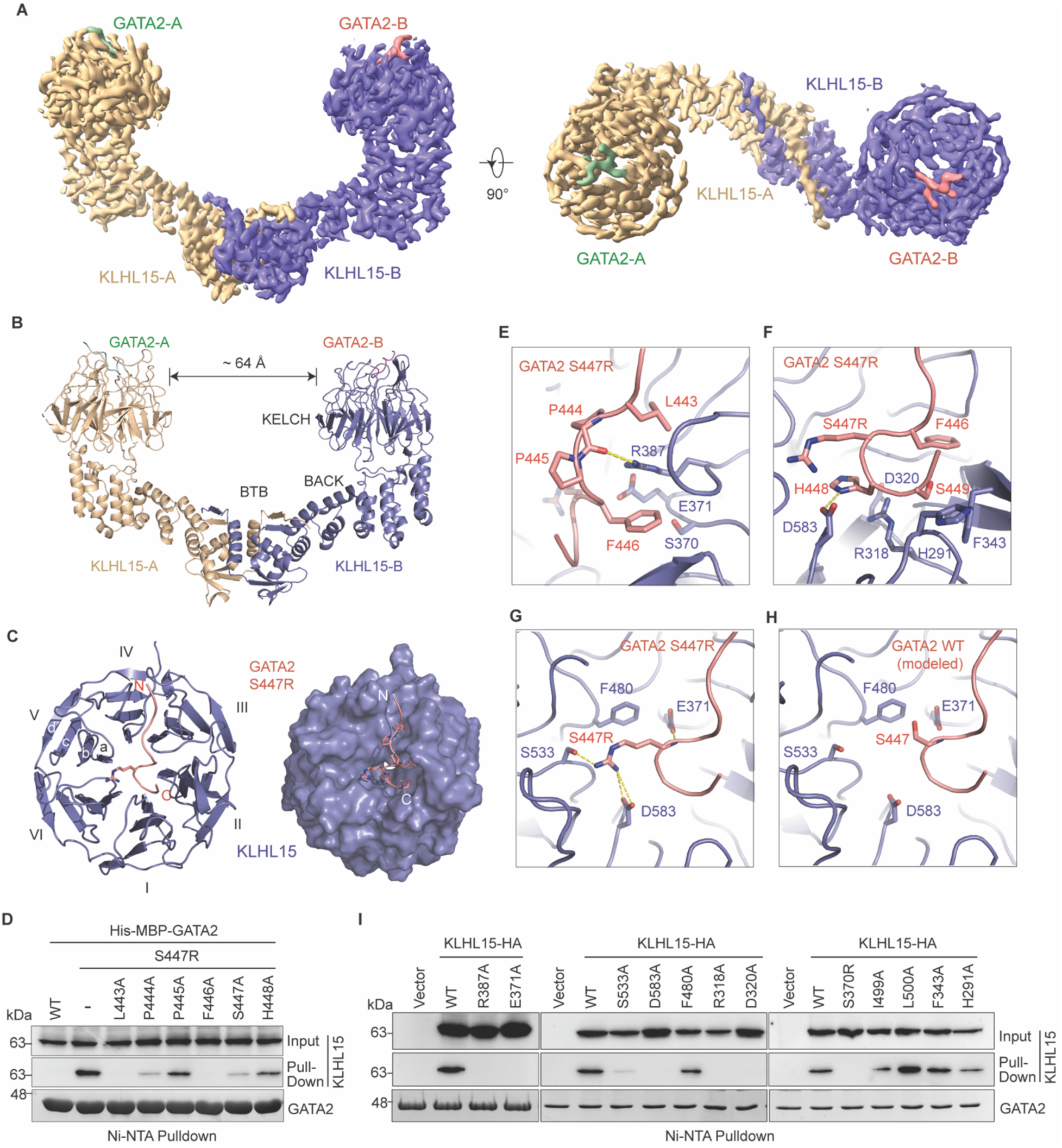
Structural mechanism of genetic glue-enabled neo-protein protein interface. (A) Cryo-EM map of KLHL15 (wheat/slate) in complex with GATA2 neo-degron peptide (green/salmon). (B) Overall structure of the KLHL15-GATA2 complex. The closest distance between the two KELCH-repeat domains is indicated at the top of the dimer. (C) Top view of the KELCH-repeat domain bound to the GATA2 neo-degron peptide. I-VI indicates the six blades of the domain; a-d indicates the four β-strand in each blade. N, N-terminus of the GATA2 peptide; C, C-terminus of the GATA2 peptide. (D) Ni-NTA pulldown using His-MBP tagged GATA2 peptides against 293T cell lysates stably expressing KLHL15-HA. (E) Interface between KLHL15 and the N-terminal half of the GATA2 neo-degron. Residues involved in interfacial interactions are shown in sticks. Yellow dotted line indicates hydrogen bond. (F) Interface between KLHL15 and the C-terminal half of GATA2 neo-degron. (G) Key interactions nucleated by the GATA2 S447R mutation. Yellow dotted line indicates hydrogen bond or salt bridges. (H) A modeled interface of GATA2 wildtype, in which Arg447 was simply mutated back to Ser447. (I) Ni-NTA pulldown using His-MBP tagged GATA2 S447R peptide against 293T cell lysates stably expressing wildtype or indicated mutants of KLHL15-HA.

KLHL15 contains an N-terminal BTB domain, a central BACK domain and a C-terminal KELCH-repeat propeller domain (**Fig. 5B**). Like many other BTB-KELCH domain proteins, KLHL15 adopts a dimeric architecture mediated via its BTB domain.^42,43^ We have also determined the structure of KLHL15 in complex with CUL3 at a global resolution of 3.7 Å **(Fig. S5A)**. Assembling with CUL3 does not change the overall architecture of KLHL15 and the relative positions of the two KELCH-repeat domains **(Fig. S5B)**.

### KLHL15 and GATA2 neo-degron interface

The KELCH-repeat domain of KLHL15 forms a canonical six-bladed propeller, with a central pocket that measures ∼19 Å in length and ∼11 Å in width. (**Fig. 5C**). The GATA2 neo-degron is anchored at the KLHL15 substrate binding pocket in a coiled conformation, primarily packing against blade II and III of the E3’s propeller (**Fig. 5C**). At the center of the neo-degron, the serine to arginine disease mutation enables the peptide to reach blade V and VI on the opposite side of the KELCH domain, locking the neo-degron in place via multivalent interactions. Together, the disease mutation site and its flanking residues constitute an extensive protein-protein interaction (PPI) interface with a total buried surface area of ∼1,500 Å^2^.

The N-terminal half of the neo-degron features two hydrophobic residues, Leu443 and Phe446 (referred to as -4 and -1 position, respectively), which are inserted into two neighboring grooves on the KLHL15 propeller (**Fig. 5E**). Mutating them into alanine (L443A and F446A) abolished the GATA2-KLHL15 interaction in an *in vitro* pull-down assay, highlighting their important roles at the interface (**Fig. 5D**). The two proline residues in between, Pro444 and Pro445, stabilize the conformation of the neo-degron mainly via backbone hydrogen bonding with KLHL15 (**Fig. 5E**). Consistently, the P444A and P445A mutations disrupt GATA2 and KLHL15 interactions to a lesser extent (**Fig. 5D**). These results are in agreement with prior saturation mutagenesis analysis of endogenous KLHL15 substrates, which shows that variations at these two positions are more tolerated than variations at the -4 and -1 sites.^19^ KLHL15 Arg387 and Glu371 are key residues that neutralize each other and shape the wall of the hydrophobic grooves housing Leu443 and Phe446 of GATA2. The guanidinium group of KLHL15 Arg387 also makes the hydrogen bonds with GATA2 backbone (**Fig. 5E**). Alanine mutations of these two residues (R387A and E371A) abrogated GATA2 binding (**Fig. 5I**). Similarly, mutating KLHL15 Ser370 into a bulkier residue (S370R), which is predicted to sterically clash with GATA2 Phe446, also abolished GATA2-KLHL15 interactions.

At the C-terminal half of the neo-degron, the FRH motif and its following Ser449 residue form a β-turn (**Fig. 5F**). Located at the tip of the β-turn, the “R” and “H” in the motif (Ser447Arg and His448) are deeply inserted into the central pocket of the E3. GATA2 His448 makes polar contacts with KLHL15 Asp583 and Arg318. A nearby aspartate residue, Asp320 neutralizes the charges of Arg318, and together they construct the base of the substrate degron recognition pocket. In agreement with their involvement in recognizing the FRH motif, KLHL15 R318A and D320A mutations abrogated GATA2-KLHL15 interaction. Consistently, H448A mutation in GATA2 also diminished the neo-degron-E3 binding (**Fig. 5D, 5I**).

Overall, the GATA2 neo-degron makes continuous contacts with KLHL15 and is constrained into a defined conformation. The numerous interactions contributed by residues flanking the disease mutation establish a sub-optimal interface between GATA2 and KLHL15, forming the basis for the mutation-enhanced neo-interface. These basal interactions are manifested by the single digit micromolar affinity of GATA2 and KLHL15 as measured in our biophysics studies.

### Genetic glue mechanism of mutation-enabled neo-PPI

As the hallmark of the GATA2 neo-degron, Arg447 is deeply embedded into the KLHL15 pocket, functioning as a key anchor at the interface. The aliphatic side chain of Arg447 packs against the phenyl ring of KLHL15 Phe480, while its guanidinium group forms a hydrogen bond with KLHL15 Ser533 and a salt bridge with KLHL15 Asp583 (**Fig. 5G**). Additionally, the amide group of the Arg447 backbone donates a hydrogen bond to the side chain of KLHL15 Glu371, further securing the neo-degron-E3 binding. Alanine mutations at these positions (KLHL15 S533A, D583A, F480A and E371A) all impacted GATA2 binding, albeit at different degrees (**Fig. 5I**). Consistent with the key interactions introduced by the serine to arginine mutation, when wildtype GATA2 Ser447 is modeled in the structure, these interactions would be mostly lost (**Fig. 5H**). Above GATA2 Arg447, we observed densities that could be traced to the long b-c loop from KLHL15 blade V (**Fig. S5C**). Due to the flexible nature of this loop and its fragmented densities, we could not confidently model its conformation. Yet, it is clear that this loop further buries Arg447, shielding it from the solvent. Therefore, the single amino acid substitution in GATA2 nucleates an intimate interface with KLHL15 via a network of hydrophobic interactions, hydrogen bonds and electrostatic interactions, enabling the high-affinity neo-degron-E3 interaction.

In summary, our study reveals the structural basis of how KLHL15 specifically recognizes the GATA2 S447R mutant as a neo-substrate. By leveraging the intrinsic weak affinity between GATA2 and KLHL15, the serine-to-arginine substitution introduces a critical bulkier side chain that physically extends the sub-par GATA2-KLHL15 interface and drives their high-affinity productive binding. The GATA2 S447R mutation, therefore, acts as a genetic glue, echoing the molecular glue mechanism highlighted by plant hormone auxin and the therapeutic agent, immuno-modulatory drugs (IMiDs).^9,44–48^

### Additional disease mutations that generate FRY-like neo-degrons

The stringent threshold (top 2.5%) applied in our initial screen of disease mutation-rendered internal degrons might have excluded additional KLHL15 neo-substrates with a slightly lower activity than the GATA2 S447R mutant. Furthermore, we only focused on pathogenic or likely pathogenic mutations in transcription factors, which represent a small fraction of missense variants deposited in ClinVar. Given the PTM-free nature and sequence simplicity of the FRY-like motif, it is conceivable that many other disease-associated mutations could generate a KLHL15 degron in functionally diverse proteins.

To identify additional FRY-like neo-degrons, we searched the missense variants in ClinVar that are located in IDRs of the entire human ORFome and identified ∼1,600 mutations that create an FRY-like motif based on its previously characterized sequence profile.^23^ We then selected a small subset of these mutations for experimental validation based on their known biological functions, protein lengths and sub-cellular locations (**Fig. S6A**). The mutant peptide sequences were cloned in the GPS system and assessed for their degron activities. Among these, several mutations stood out with a strong destabilization signal, including the S430R mutant of mRNA decay activator protein, ZFP36L2, and the S155F mutant of mitogen-activated protein kinase kinase kinase 20 (MAP3K20) (**Fig. 6A**). We also identified mutations that induce a moderate shift in the GPS assay, exemplified by plakophilin-2 (PKP2) P192L, and mitotic checkpoint serine/threonine-protein kinase BUB1 beta (BUB1B) N598Y. When tested in co-immunoprecipitation experiments, these peptides indeed showed association with KLHL15, suggesting that their activities are dependent on KLHL15 (**Fig. S6B**). Interestingly, some peptides bearing a mutation-created FRY-like motif showed little, if any, degradation activity or KLHL15 interaction (**Fig. S6A**). Therefore, a FRY-like tripeptide motif is insufficient to confer high affinity towards KLHL15. Although alignment of KLHL15 neo-substrates does not reveal a consensus sequence beyond the central FRY-like motif (**Fig. 6B**), we find that charged amino acids are generally disfavored in its immediate flanking regions, consistent with previously reported deep mutagenesis scan results for several known FRY degrons (**Fig. S6B, S6C**)^19^. A comprehensive investigation on the flanking sequences is warranted to define the complete degron motif recognized by the E3.

**Figure 6.**
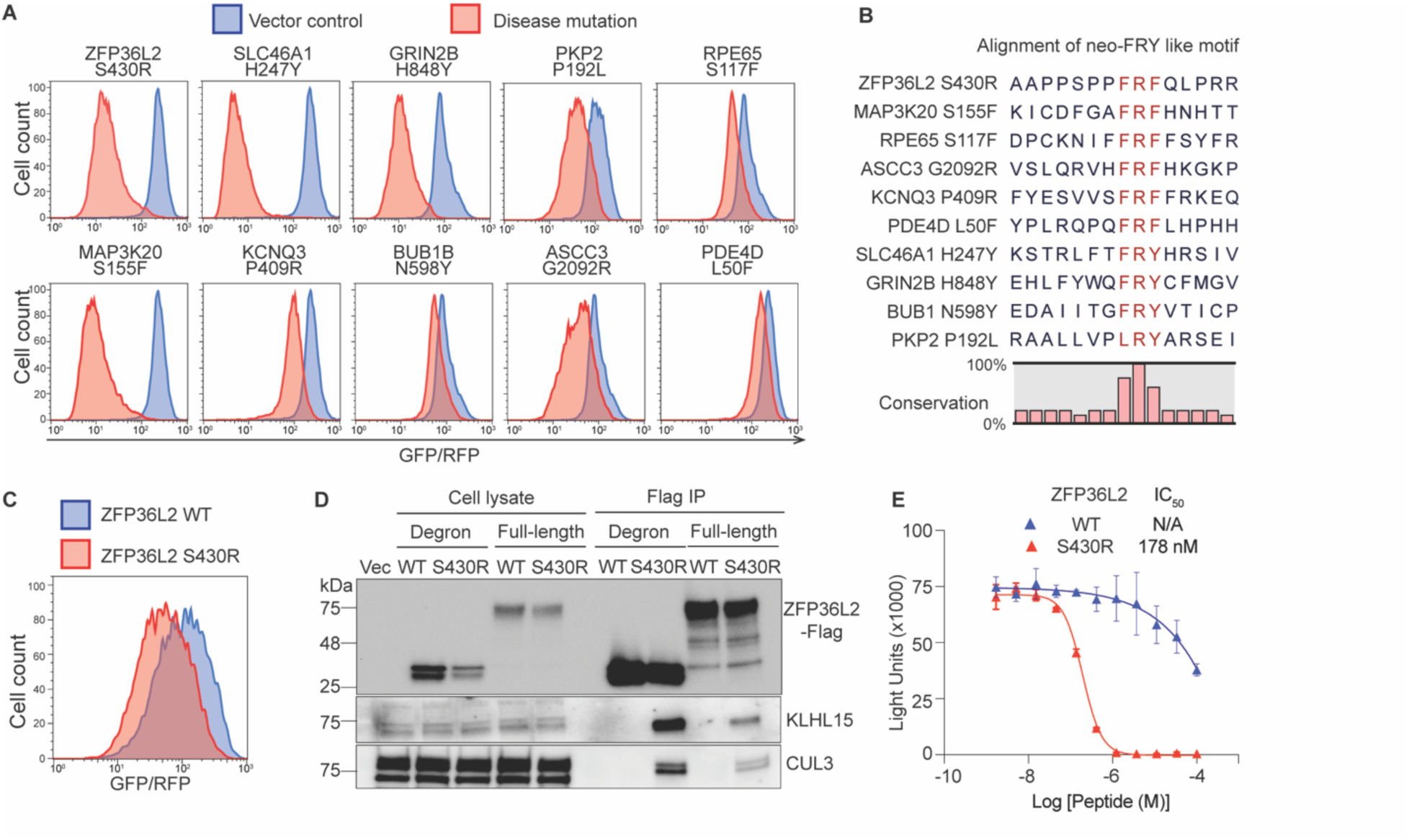
Additional disease mutations that generate FRY-like neo-degrons. (A) FACS analysis of degron activities in peptide-based reported assays. (B) Alignment of neo-FRY like motifs that show degradation activity. (C) FACS analysis of full-length ZFP36L2 protein stability. (D) Immunoblots of Flag IP from 293T cells expressing ZFP36L2 degrons or full-length proteins. (E) AlphaLISA competition assays for assessing ZFP36L2 peptides binding to KLHL15. Data were presented as mean ± s.d. n = 3 biologically independent samples. IC_50_, half-maximum inhibitory concentration. N/A, not available.

One of the functional neo-degrons validated by the GPS assay belongs to ZFP36L2, whose loss of function leads to female infertility, cardiomyopathy, and hematopoiesis deficiency.^49–51^ To further confirm that genetic mutations create KLHL15 substrates, we cloned the wildtype and mutant form of full-length ZFP36L2 protein and found that ZFP36L2 S430R indeed showed lower steady levels (**Fig. 6C**). Moreover, wildtype ZFP36L2 did not co-precipitates with KLHL15, while ZFP36L2 S430R mutant pulled down KLHL15 and CUL3 (**Fig. 6D**). Finally, we quantified the binding of ZFP36L2 degron peptides to KLHL15 in an AlphaLISA assay. In contrast to the wildtype ZFP36L2 peptide, which showed competition at ∼ 100 μM, the S430R mutant exhibited an IC_50_ of ∼180 nM (**Fig. 6E**). Together, our results demonstrate that ZFP36L2 S430R mutant represents a neo-substrate of KLHL15.

## DISCUSSION

Although genomic testing has greatly facilitated the detection of genetic variants, most variants identified to date remain insufficiently characterized and therefore cannot inform clinical diagnosis or treatment. In this study, we have developed a novel tricolor reporter system, iGPS, which allows us to both map *bona fide* internal SLiM degrons at scale and assess the impact of disease mutations within the non-structured regions of the human proteome on protein stability. By profiling the disease-associated variants in the IDRs of transcription factors, we identified mutations that unexpectedly induce specific neomorphic protein interactions, leading to a gain-of-degron and loss-of-function in the protein. Unlike mutations in protein domains that compromise protein folding and evoke protein quality control mechanism for clearance, the IDR mutations in GATA2 and ZFP36L2 give rise to a functional SLiM-based degron that is recognized by a specific E3, KLHL15. It is conceivable that additional protein-destabilizing IDR mutations uncovered in our studies may act through similar mechanisms, warranting further investigation to identify the responsible E3s. Development of inhibitors against these E3s, such as KLHL15, might represent a viable approach to treat the corresponding genetic diseases. Such a therapeutic strategy, nevertheless, is at the risk of interfering with the normal functions of the relevant E3s. Overall, our findings highlight a previously underappreciated tradeoff inherent to the UPS, the same degron diversity regulating proteostasis also invites mutation-induced neo-degron formation and instability.

Importantly, our mechanistic studies of the neomorphic GATA2-KLHL15 interaction help establish “genetic glue” as a new concept in induced proximity. We propose to use this term to describe general mutations at a PPI interface that introduce extra chemical moieties, either as part of an amino acid side chain or via amino acid insertions in the context of a protein fold, to complement an otherwise sub-optimal protein contact site and create a specific neo-interaction. Such a mechanism of action is reminiscent of PTM-mediated PPIs, in which amino acid modifications, such as phosphorylation, hydroxylation, methylation and acetylation, effectively extend PPI interface and augment sub-part binding between two proteins.^4^ In both scenarios, the added moieties rely on the intrinsic weak or moderate interactions made by their flanking regions to achieve high affinity specific binding. Analogous to the effect of the single amino acid S447R mutation in GATA2, the latent HIF1α degron is functionalized by the hydroxylation of its central proline residue that enhances its affinity to VHL by ∼ 1,000 folds, from 20 μM to 20 _nM._52,53

Noticeably, this same principle is shared by molecular glues, which act at a loosely packed PPI interface to stabilize the ternary interaction with high cooperativity.^9^ Despite the covalent versus non-covalent nature of genetic and molecular glues, their common mechanism of action suggests that they might be interchangeable for the same PPI interface. In fact, recent studies have shown that the medulloblastoma insertion mutations of KBTBD4 can be structurally and functionally mimicked by UM171, a potent agonist of haemopoietic stem cell *ex vivo* expansion, to promote its neo-interactions with HDAC1/2.^17,54^ Such a converging mechanism between genetic and molecular glues is paralleled by the functional replacement of a phosphorylation site within the β-catenin degron by a synthetic molecular glue compound for high affinity binding to the β-TrCP E3 ligase.^55^ Therefore, screening and identifying genetic glues that create neo-degrons in therapeutic targets could inform prospective discovery of novel molecular glue degraders.

Although the current study focuses on the FRY-like motifs recognized by KLHL15, a variety of PTM-independent internal degrons have been identified for other E3s, such as the [DE]xE[TS]GE motif recognized by KEAP1 and the [KR]RxL[DE] motif recognized by CCNF.^10,56,57^ Using the same workflow we applied to search for latent FRY-like motifs, we have identified 25 and 256 disease mutations in ClinVar that generate motifs matching those recognized by KEAP1 and CCNF, respectively. Future studies will help validate whether these mutation-created motifs are sufficient to trigger protein degradation by KEAP1 and CCNF1. Besides degrons, a variety of SLiMs, such as the LxCxE motif recognized by retinoblastoma (Rb) protein and the [WFY]xx[LVI] LC3-interacting region (LIR) motif in autophagy,^58,59^ are characterized by multiple bulky amino acids and known to mediate PPI in diverse cellular pathways. Single amino acid substitution in otherwise unrelated proteins might be sufficient to create these SLiMs.^60^ Beyond SLiMs from IDRs, amino acid insertions at the surface of a folded protein domain can also render genetic glues to promote neo-PPIs, as evidenced by the KBTBD4 mutations. Genetic glues, therefore, not only represent a recurring mechanism in disease mutations that rewires protein network, but also afford a potential strategy to guide molecular glue discovery for novel therapeutics development through induced proximity.

**Figure S1.**
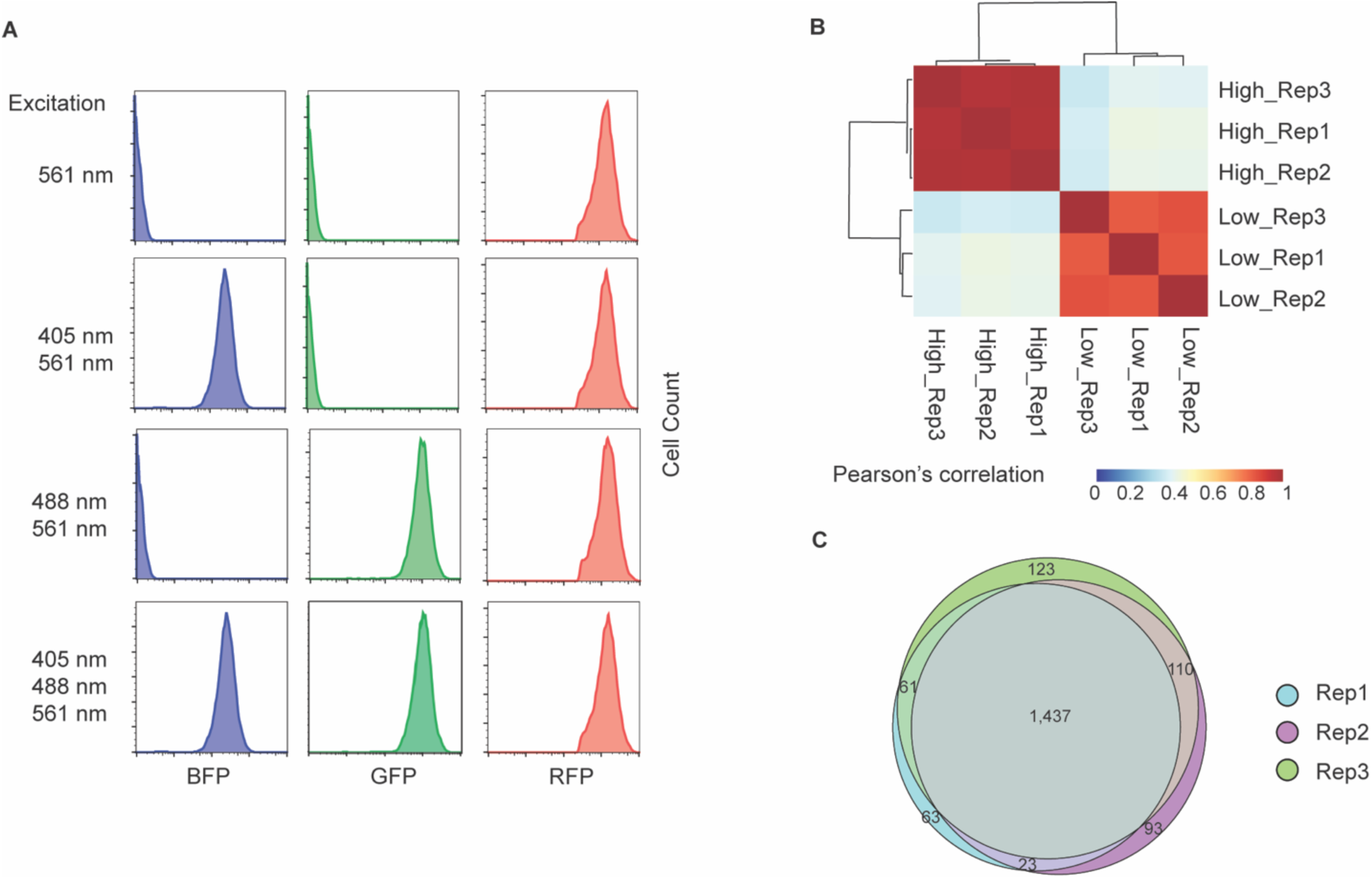
Systematic analysis of the effects of disease-associated IDR mutations on transcription factor stability. (A) FACS analysis of HEK293T cells expressing the iGPS reporter under varying excitation laser configurations. Each row corresponds to a distinct excitation condition. BFP, GFP and RFP are excited using lasers at 405 nm, 488 nm and 561 nm, respectively. (B) Hierarchical clustering of samples from three independent biological replicates based on normalized oligo abundance profiles derived from FACS-sorted low- and high-stability populations. Heat map values indicate pairwise correlations between samples, demonstrating reproducibility across replicates and clear separation between stability groups. (C) Overlap of degradation-promoting peptides (log₂FC ≥ 3) identified across three independent biological replicate screens.

**Figure S2.**
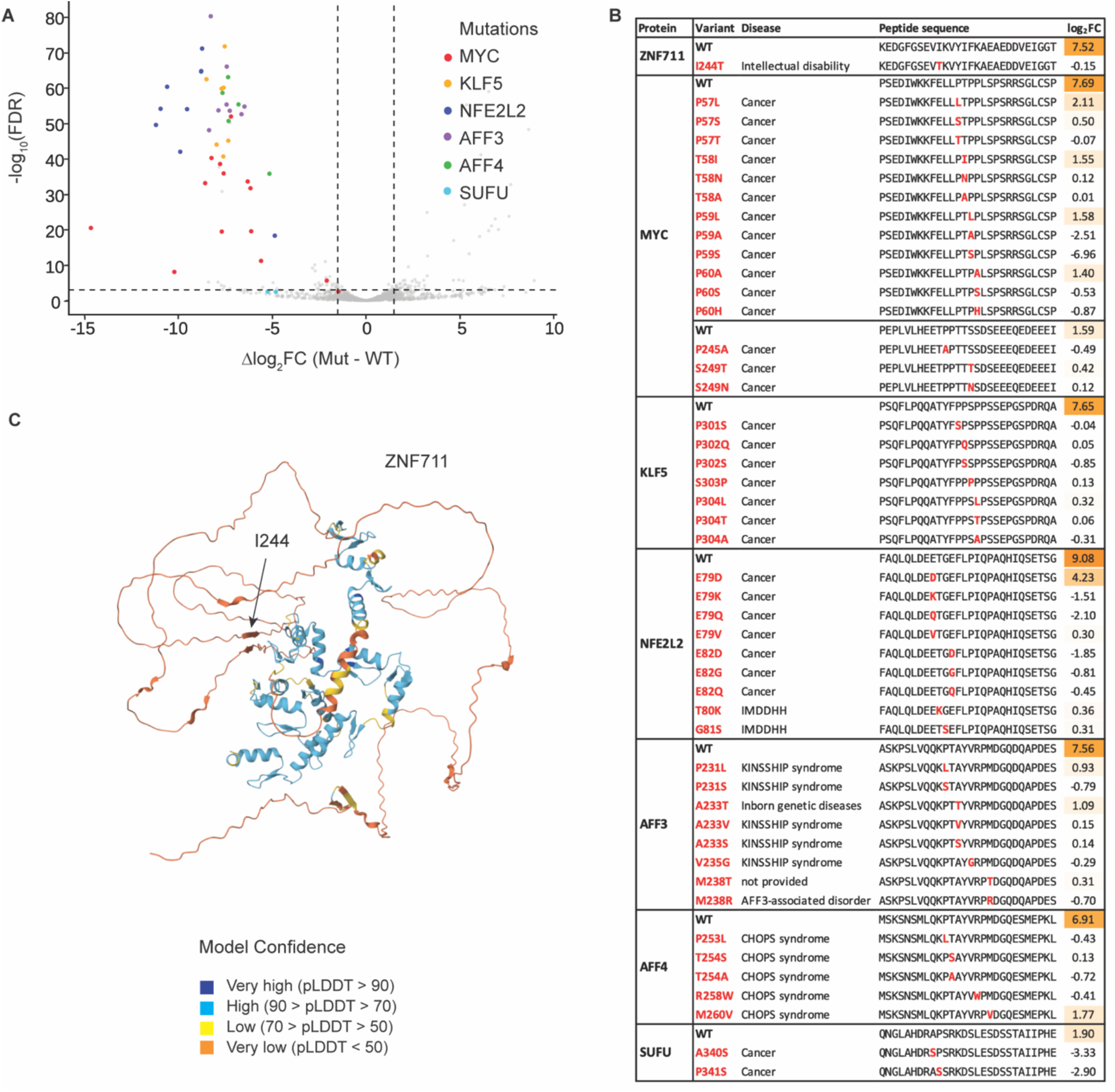
Inhibition of degron activity by disease-associated mutations in transcription factors. (A) Volcano plot showing previously reported mutations that suppress degron activity. Known mutations are highlighted in color, whereas all other variants are shown in grey. Dashed lines indicate thresholds for statistical significance (FDR < 0.001) and effect size (|Δlog₂FC| ≥ 1.5). (B) Representative degron-disrupting variants. The table lists representative variants that suppress degron activity, together with their associated diseases, log₂FC values, and peptide sequences. Mutated residues are highlighted in red. Degron activity is represented by log₂FC values, with color intensity indicating degradation strength. (C) AlphaFold-predicted structure of ZNF711 with the mutated residue I244 indicated.

**Figure S3.**
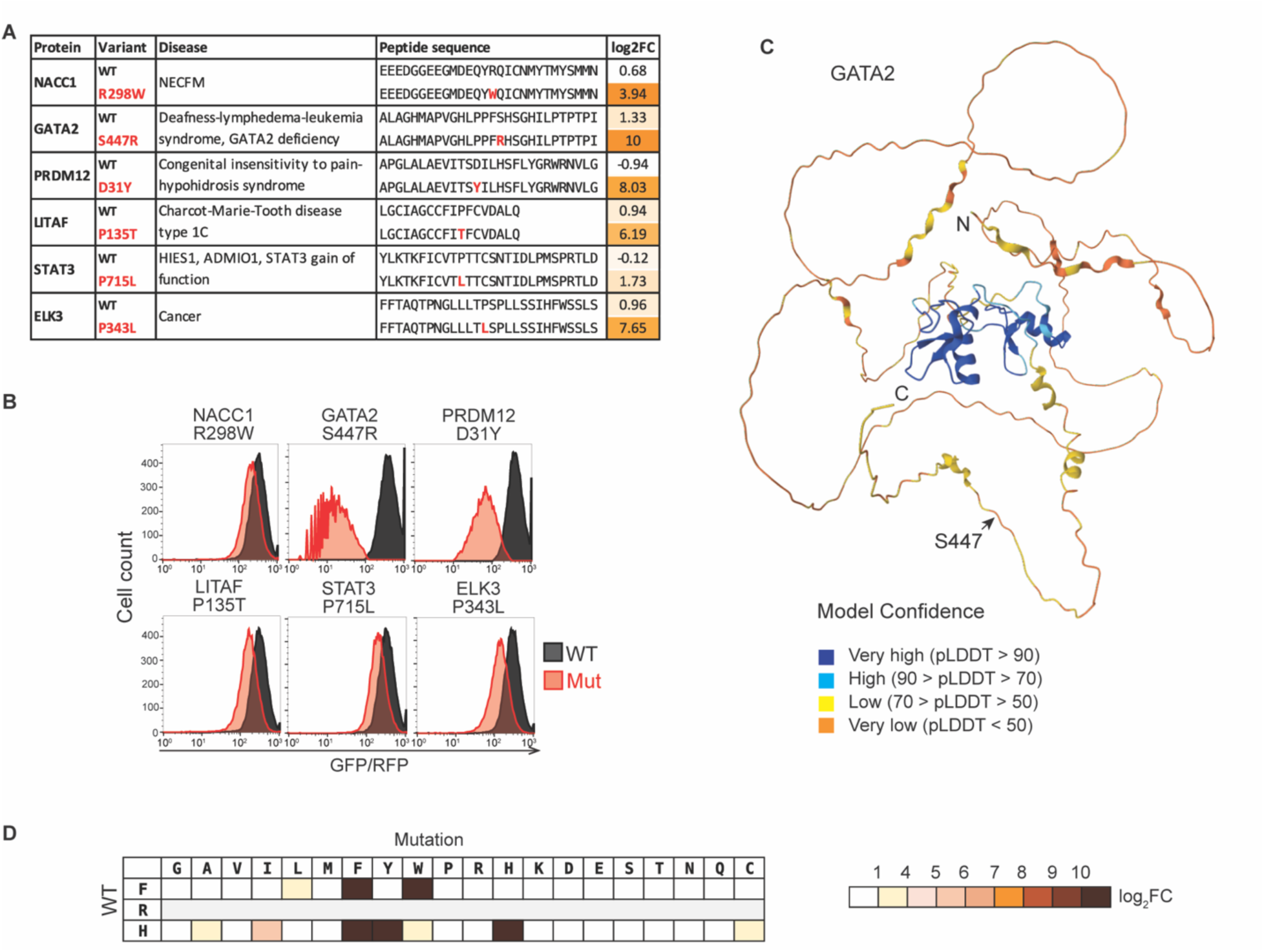
Neodegron formation driven by disease-associated mutations in transcription factors. (A) Table listing representative variants that generate neodegrons. Associated diseases, log₂FC values and peptide sequences are provided, with mutated residues highlighted in red. (B) Validation of mutation-induced protein degradation using a peptide-based iGPS reporter assay. (C) AlphaFold-predicted structure of GATA2 with the S447R disease-associated mutation site indicated together with the color scheme of the pLDDT scores. (D) Mutagenesis analysis of the GATA2 FRH degron motif. Degron activity is shown as log₂FC, with color intensity reflecting degradation strength.

**Figure S4.**
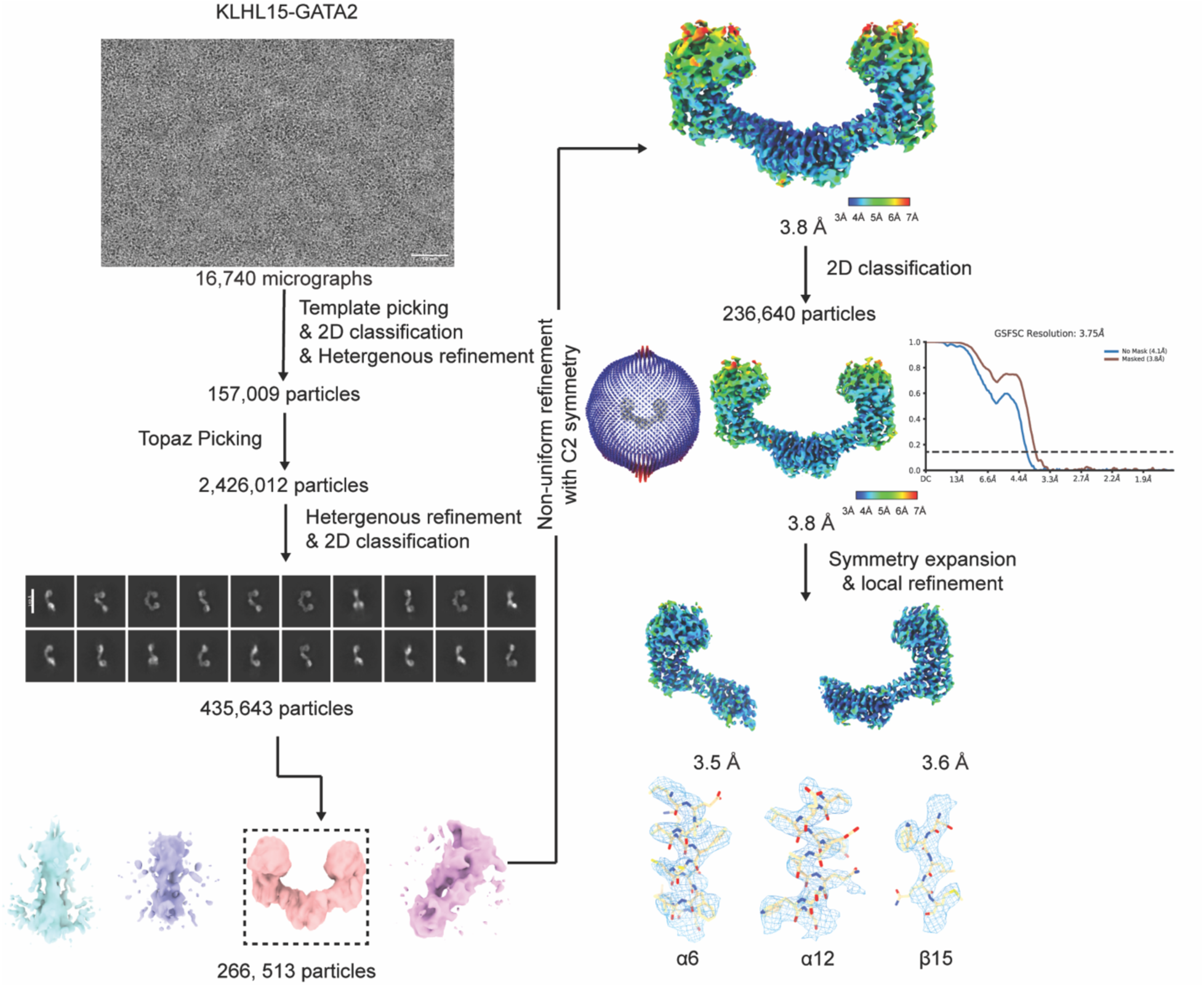
Cryo-EM data processing for the KLHL15-GATA2 complex. A representative cryo-EM micrograph, typical 2D classifications, local resolution maps, Fourier shell correlation (FSC) curves and density maps of representative regions of the complex were shown.

**Figure S5.**
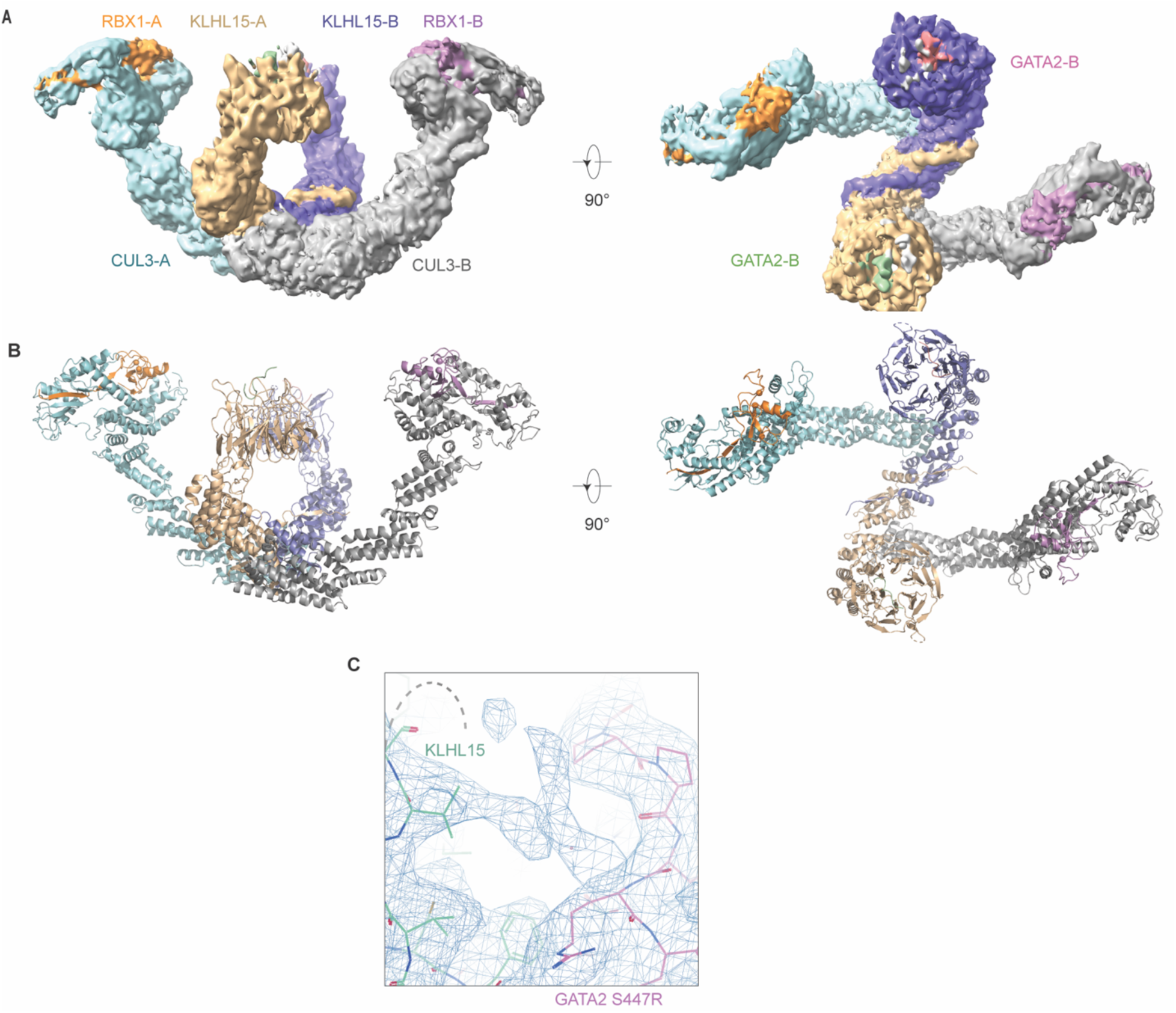
Structure of the CRL3^KLHL15^-GATA2 complex. (A) Cryo-EM map of the CRL3^KLHL15^-GATA2 complex. CUL3 (cyan/grey), RBX1 (orange/violet), KLHL15 (wheat/slate), GATA2 (green/salmon). (B) Overall structure of the CRL3^KLHL15^-GATA2 complex. (C) Density of the b-c loop from KLHL15 blade V.

**Figure S6.**
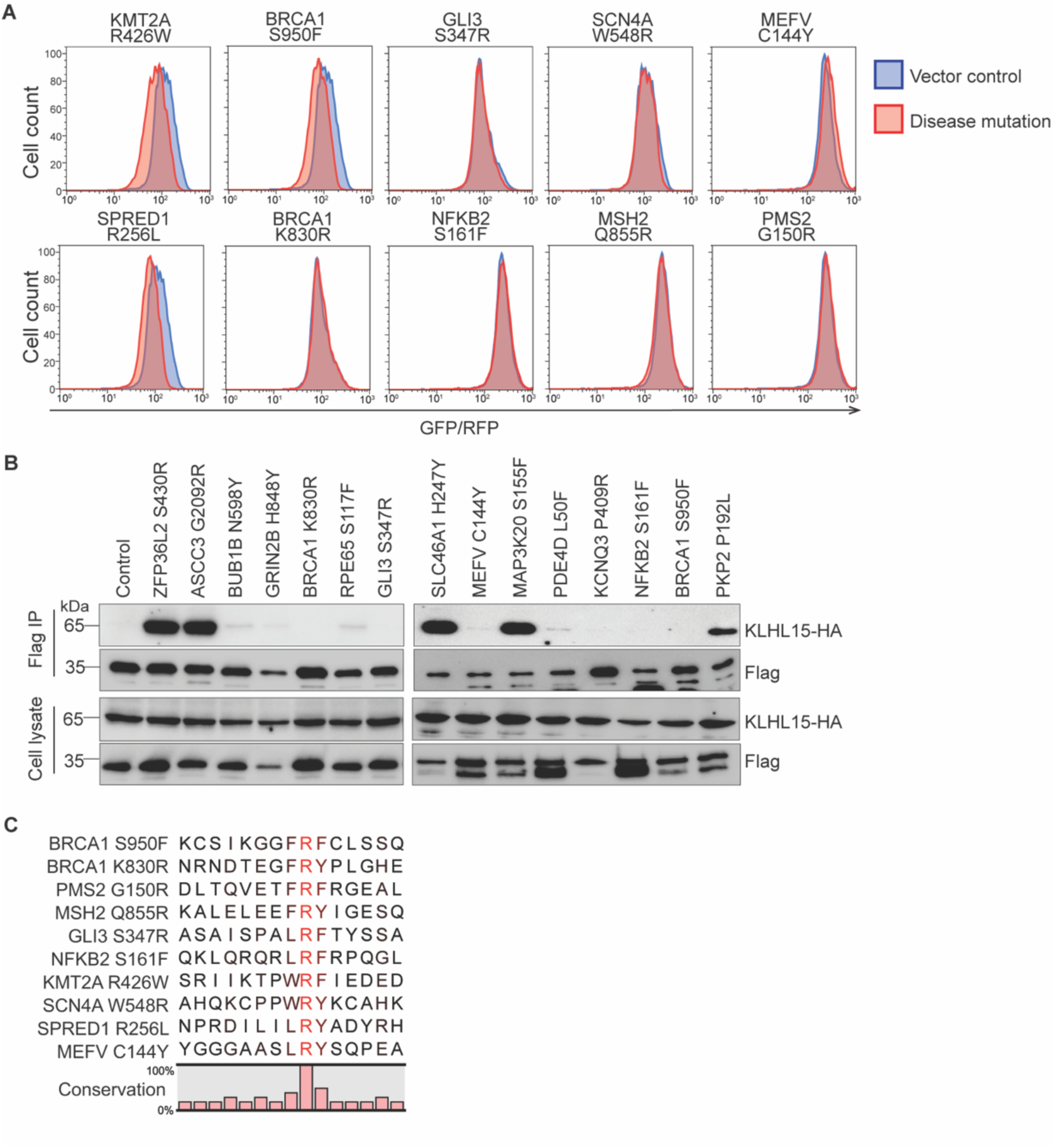
Disease mutations create additional FRY-like neo-degrons. (A) Degron activities of disease-mutation generated FRY-like motifs were evaluated in peptide-based FACS reported assays. (B) Flag IP immunoblot analysis of 293T cells expressing different FRY-like peptides. (C) Alignment of FRY-containing motifs with negligible or no degradation activity.

## METHODS

### Cell culture and drug treatments

HEK293T cells (ATCC CRL-3216) were maintained in DMEM (Gibco) supplemented with 10% fetal bovine serum (HyClone), 100 U ml⁻¹ penicillin, and 100 μg ml⁻¹ streptomycin at 37 °C in a humidified incubator with 6% CO₂. ExpiSf9 cells (ThermoFisher Scientific, A38841) were cultured according to the manufacturer’s instructions.

Proteasomes were inhibited with bortezomib (1 μM, 5 h; BioVision) or MG-132 (10 μM, 5 h; Merck Millipore). E1 activity was inhibited with TAK-243 (1 μM, 5 h; MedChemExpress). Cullin-RING ligase (CRL) activity was suppressed by treatment with MLN4924 (1 μM, 5 h; Active Biochem) or by transduction with dominant-negative Cullin (DNCUL) lentivirus (multiplicity of infection (MOI) ∼10, 40 h). IKZF degron-mediated degradation was induced with pomalidomide (1 μM, 6 h; Selleckchem). For CHX-chase assays, cells were treated with cycloheximide (100 μg ml⁻¹; Calbiochem), and samples were collected at the indicated time points.

### Lentivirus production

Lentiviral particles were produced by co-transfecting HEK293T cells (∼80% confluency) with pRev, pTat, pHIV gag/pol, pVSV-G, and the lentiviral transfer plasmid using TransIT-293 (Mirus Bio) according to the manufacturer’s instructions. The culture medium was replaced 24 h post-transfection, and viral supernatants were harvested 48 h later.

### Plasmid construction

The iGPS lentiviral backbone (pLenti-iGPS) was constructed by inserting an in-frame TagBFP–SGFP2 fusion linked by a glycine–serine linker (SGSGSSAGSGSG), followed by an IRES–DsRed cassette, into a pLenti vector under the control of the EF1α promoter. A 20-amino-acid C-terminal cap (QGRARPNQEVQIGEMENQLS), used to suppress artificial C-degrons in the pLenti-GPS reporter, was adopted from a previous study^19^.

iGPS and GPS reporter constructs were generated by cloning peptides or genes of interest into the pLenti-iGPS and pLenti-GPS vectors using Gibson assembly (New England Biolabs) or Gateway recombination (Invitrogen), respectively, according to the manufacturers’ instructions.

The open reading frame (ORF) of human KLHL15 and ZFP36L2 was purchased from Horizon Discovery. For protein expression in insect cells, KLHL15 cDNA was cloned into pACE vector with a N-terminal His-Venus tag. For mammalian expression, full-length KLHL15 was cloned into a pLC lentiviral vector (Addgene #123322). Full-length GATA2 and ZFP36L2 were cloned into the pLenti-GPS vector with a C-terminal GFP tag.

### iGPS/GPS assays

iGPS and GPS reporter cell lines were generated by lentiviral transduction at a low MOI of ∼0.2, followed by puromycin selection (1 μg ml⁻¹; Clontech). GFP/RFP and BFP/RFP fluorescence ratios were quantified by flow cytometry using a BD LSRFortessa system (BD Biosciences). TagBFP (BFP), SGFP2 (GFP) and DsRed (RFP) were excited with 405-nm, 488-nm and 561-nm lasers, respectively. Data were acquired at a rate of 5 × 10³–1 × 10⁴ cells s⁻¹. Cells were gated by doublet discrimination and DsRed positivity to identify single reporter-expressing cells. At least 10,000 single cells were analyzed per sample. No fluorescence compensation or background subtraction was applied. Flow cytometry data were analyzed using FlowJo.

The peptide sequences used to evaluate iGPS performance were as follows: IKZF (RPFQCNQCGASFTQKGNLLRHIKLH); MYC (PSEDIWKKFELLPTPPLSPSRRSGLCSP); MYC T58A (PSEDIWKKFELLPAPPLSPSRRSGLCSP); NFE2L2 (FAQLQLDEETGEFLPIQPAQHIQSETSG); NFE2L2 T80K (FAQLQLDEEKGEFLPIQPAQHIQSETSG); USP1 (VSIRVGGNTQPSKVLNKKNVEAIGLLGG); ARSB (RHDLSREYPHIVTKLLSRLQFYHKHSVP); and BRWD1 (STFSTAFPGTMYQHIKMHRRILGHLSAV).

### Comparison of the iGPS and GPS systems

For comparison of the iGPS–GPS systems, an oligonucleotide library was designed from IDRs of 424 cancer-related genes with predicted AlphaFold pLDDT scores <85. Regions longer than 28 residues were tiled into overlapping 28-residue peptides with 10 residues of overlap, generating a library of 5,989 oligonucleotides. The oligonucleotide pool (Twist Bioscience) was PCR-amplified using KAPA HiFi HotStart DNA Polymerase (Roche) and cloned into the pLenti-iGPS and pLenti-GPS vectors by Gibson assembly at approximately 500-fold coverage per-oligonucleotide.

To generate iGPS and GPS cell libraries, HEK293T cells were transduced with the corresponding lentiviral libraries at an MOI of 0.1, providing approximately 500-fold coverage per oligonucleotide. Cells were selected with puromycin (1 μg ml⁻¹) for 7 days. iGPS and GPS cell libraries, together with vector-only controls, were analyzed by flow cytometry, with 5 × 10⁵ cells acquired per sample.

### iGPS screening of disease-associated IDR missense mutations in transcription factors

Germline variants annotated as pathogenic or likely pathogenic in 1,953 human transcription factors (TFs) were collected from ClinVar (May 2023 release), and somatic cancer mutations in 135 TFs annotated as oncogenes or tumor suppressors were obtained from COSMIC (May 2023 release). Variants predicted to reside within structured regions (pLDDT ≥85; AlphaFold) were excluded, yielding a total of 2,081 IDR-localized missense mutations.

A pooled oligonucleotide library encoding wild-type–mutant peptide pairs (10–28-amino acids) was synthesized (Twist Bioscience), PCR-amplified using KAPA HiFi HotStart DNA polymerase, and cloned into the pLenti-iGPS vector by Gibson assembly at approximately 1000-fold coverage per oligonucleotide. Whenever possible, the mutated residue was positioned within the central 10 amino acids of each peptide.

Lentiviral iGPS libraries were generated and used to transduce HEK293T cells at a low MOI of ∼0.05, achieving approximately 1,000-fold cellular coverage. Following puromycin selection for 10 days, DsRed-positive cells were sorted into low-(bottom 2.5%) and high-stability (top 90%) populations based on BFP/RFP and GFP/RFP fluorescence ratios using a BD FACSAria III cell sorter. Cells were sorted twice to improve purity, and the entire screen was performed in three independent biological replicates. Approximately 2 × 10⁶ and 4 × 10⁷ cells were collected for the low- and high-stability fractions, respectively, per replicate. Genomic DNA was isolated from sorted cells, and integrated oligonucleotides were PCR-amplified using Super-Run EX Taq DNA polymerase (PRO TECH) before Illumina sequencing.

### Data processing and statistical analysis of iGPS screening

The TF IDR disease-variant screen was performed together with several additional iGPS screens unrelated to this study, and all sequencing data were processed jointly using a common analysis pipeline. Oligonucleotides with fewer than 50 total sequencing reads across the six samples (Low_rep1, High_rep1, Low_rep2, High_rep2, Low_rep3, High_rep3) were excluded from downstream analyses.

Differential enrichment between low- and high-stability fractions was analyzed in R using DESeq2, which fits a negative binomial generalized linear model with library-size normalization and dispersion estimation. To account for variability among screening replicates, the screening replicate was included as a blocking factor in the design formula (∼ screen + bin). Effect sizes were reported as log₂ fold changes (log₂FC) comparing the low- and high-stability fractions. Statistical significance, including one-tailed tests for enrichment in the low-stability fraction (padj_one_tailed), was assessed using the Wald test with Benjamini–Hochberg false discovery rate (FDR) correction.

For each variant, the mutation effect was defined as Δlog_2_FC = log _2_ FC_Mut_ − log _2_ FC_WT_. Statistical significance of Δlog_2_FC was evaluated using a two-sided z-test under the standard normal distribution, and *P* values were adjusted for multiple testing using the Benjamini–Hochberg procedure. Variants were considered significant if the FDR was < 0.001 and |Δlog₂FC| ≥ 1.5.

### Degron mapping and characterization

Minimal degrons were mapped by serial truncation, alanine scanning, and saturation mutagenesis using a pooled iGPS peptide screening approach, as described above for the TF IDR disease-variant screen.

### CRISPR-Cas9 screen for E3 ligase identification

HEK293T cells stably expressing Cas9 and GPS reporters carrying full-length ZNF711 or GATA2 S447R were transduced in a 96-well array format with lentiviruses expressing sgRNAs targeting 52 CRL5 substrate receptors or 113 CRL3 substrate receptors, respectively. sgRNAs targeting CRL3 and CRL5 substrate receptors (2-4 sgRNAs per gene) were cherry-picked from the Sanger QuickPick™ Knockout gRNA library (Sigma-Aldrich) and the LentiArray™ Human Ubiquitin CRISPR Library (Thermo Fisher Scientific; cat. no. A42270). GFP/RFP fluorescence ratios were measured 120 h after transduction using a BD LSRFortessa flow cytometer (BD Biosciences). GFP/RFP ratios were normalized to all wells on the corresponding plate and converted to Z scores. Genes were ranked according to their Z scores, and the highest-ranking candidate E3 ligases were selected for validation.

The target sequences and corresponding sgRNA identifiers used in Figs. 2J and 3H are provided below:

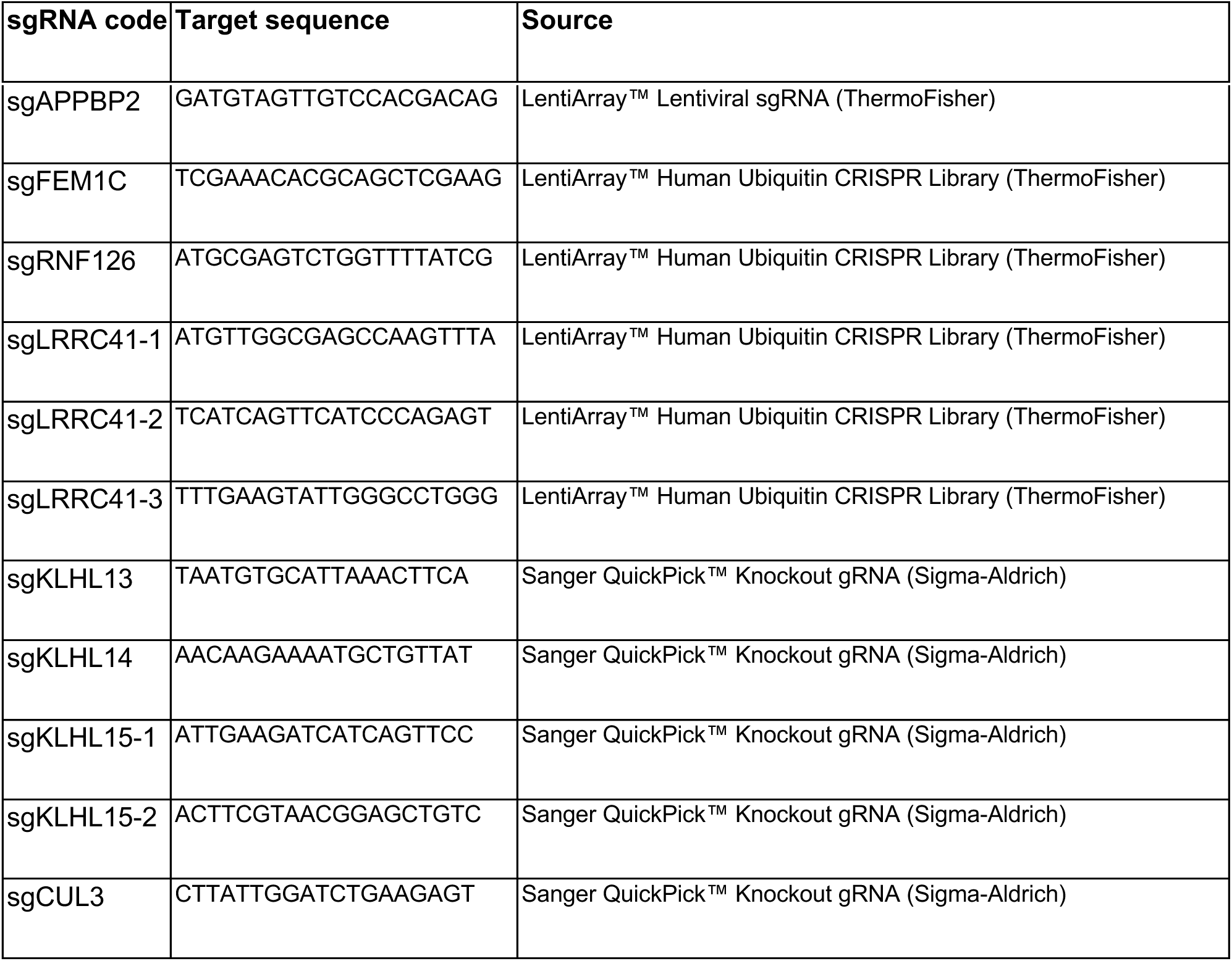

### Co-immunoprecipitation

293T cells stably expressing KLHL15 were transfected with 2 ug GATA2 (or other FRY-motif containing proteins) plasmids using FuGENE HD transfection reagent (Promega # E2311). Two days after transfection, cells were washed twice with cold PBS and lysed on ice with lysis buffer for 20 mins (20 mM Tris-HCl pH 7.5, 150 mM NaCl, 1mM MgCl2, 0.5 mM EDTA, 10% glycerol, 0.1% NP-40, protease inhibitor (ThermoFisher Scientific, #A32963). The lysates were centrifuged at 14,000 rpm for 10 mins, and the supernatants were incubated with pre-equilibrated anti-Flag magnetic beads (ThermoFisher Scientific, #A36797) for 30 mins in the cold room. Then, the beads were washed with lysis buffer 3 times and eluted with 1.5 mg/mL 3XFlag peptide (ThermoFisher Scientific, #A36805). Inputs and eluates were analyzed with SDS-PAGE gels followed by immunoblotting.

### Protein purification

Recombinant KLHL15 protein was expressed and purified from ExpiSf9 cells. Baculovirus was generated and amplified following the manufacture’s protocol. For expression, ExpiSf9 cells were grown to a density of 3 x 10^6^ cells per ml and supplemented with ExpiSf Enhancer for 16 hours before infection. 72 hours after infection, cells were harvested and washed with PBS. Cell pellets were flash frozen in liquid nitrogen for further purification. Cell pellets were resuspended in lysis buffer (20 mM Tris-HCl pH 8.0, 200 mM NaCl, 20 mM imidazole, 0.2 mM TCEP supplemented with 1 mM PMSF, 10 µM leupeptin, 0.5 µM aproptinin and 1 µM pepstatin A) and sonicated. The lysate was centrifuge at 17,500 rpm for 45 mins twice, and the supernatant were incubated with Ni-NTA resin (ThermoFisher Scientific, # 88222) for 2 hours in the cold room. Then resin was washed with 20 column volume of lysis buffer and eluted with the elution buffer (20 mM Tris-HCl pH 8.0, 200 mM NaCl, 200 mM imidazole, 0.2 mM TCEP). The eluate was incubated with tobacco etch virus protease (TEV) overnight to remove the N-terminal His-Venus tag. For proteins used in pull-down experiments, the TEV cleavage step was skipped. Then nickel eluates were further separated on an anion exchange (Cytiva 1mL Q HP) column with a 3-50% gradient elution, and peak fractions were collected and further purified by on a size exclusion column (Superdex 200 10/300 GL increase).

His-MBP tagged GATA2 degrons were expressed and purified from *E. coli* following the Ni-NTA affinity purification procedure as described above. The eluates were buffer exchanged into the storage buffer (20 mM Tris-HCl pH 8.0, 200 mM NaCl, 0.2 mM TCEP) using a desalting column (Cytiva, PD midiTrap G-25). For pull-down assays in which MBP-GATA2 degrons were used as prey, the His-tag was removed by thrombin cleavage and further purified on a size exclusion column.

The Cul3-Rbx1 complex was purified as described earlier^61^ and mixed with KLHL15 and GATA2 peptide for 30 mins on ice. Then, the CRL3-KLHL15-GATA2 complex was separated on a Superose 6 10/300 GL column. Peak fractions were pooled together and concentrated to 1.5 mg/mL and flash frozen.

### Affinity pull-down assay

Semi in vitro pull-down assay. Recombinant His-MBP tagged GATA2 degrons (WT or mutants) were diluted in the pull-down buffer (20 mM Tris-HCl pH 7.5, 150 mM NaCl, 20 mM imidazole, 10% glycerol, 0.1% NP-40) and incubated with Ni-NTA magnetic beads (ThermoFisher Scientific, #78605) for 30 mins in the cold room. The beads were then washed twice and incubated with cell lysates from 293T cells stably expressing wildtype or mutant KHL15. After 30 mins of incubation, the beads were washed three times with the pull-down buffer and then eluted with Ni-NTA elution buffer (20 mM Tris-HCl pH 7.5, 150 mM NaCl, 200 mM imidazole, 10% glycerol, 0.1% NP-40). GATA2 degrons were detected via Coomassie stain, and KLHL15 was detected with Western blot.

Ni-NTA in vitro pull-down assay. Recombinant His-venus tagged KLHL15 protein was captured by Ni-NTA magnetic beads for 30 mins in the cold room. The beads were then washed twice and incubated with MBP-tagged wildtype or S447R mutant GATA2 degrons for 30 mins. After three washes, the beads were eluted with Ni-NTA elution buffer. Inputs and eluates were separated on SDS-PAGE gels followed by Coomassie stain.

### BioLayer interferometry (BLI) measurement

The BLI experiment was carried out using the Octet Red 96 at 30C. The assay buffer contains 20 mM Tris-HCl pH8.0, 200 mM NaCl, and 0.05% BSA, 0.02% Tween 20, 0.5 mM TCEP. Streptavidin biosensors (Sartorius, #18-5019) were loaded with 200 nM biotinylated GATA2 WT or mutant peptide (439-454) and quenched with 200 nM biocytin prior to kinetic binding analysis. Different concentrations of recombinant full-length KLHL15 proteins were used as analyte. The results were analyzed by the Octet data analysis software. The association and dissociation curves were globally fit with a 1:1 ligand model. The kon and kdis values were used to calculate the dissociation constant, Kd.

### AlphaLISA experiment

AlphaLISA experiments were performed in white half-area 96 well plates (Revvity, #6002290) in a 100 uL reaction volume. The assay buffer is the same as the BLI buffer. 40 nM biotinylated CtIP peptide were diluted into the assay buffer and aliquoted in the assay plate. Nonbiotinylated GATA2 wildtype or S447R degron peptides were then added into the reaction at indicated concentrations. 2 nM recombinant His-KLHL15 protein were aliquoted into the plate, and the mixture were incubated at room temperature for 1 hour. Then, anti-His AlphaLISA acceptor beads (Revvity, #AL178C) were diluted and added to the plate followed by 30 min incubation. Finally, the streptavidin-coated AlphaScreen donor beads (Revvity, # 6760002S) were added into the mixture and incubated for an hour in dark. The Alpha signals were acquired on a Perkin Elmer microplate reader.

### Cryo-EM sample preparation and data collection

For grid preparation of KLHL15-GATA2 and CUL3-RBX1-KLHL15-GATA2, holey gold grids (Quantifoil R1.2/1.3, 300 mesh) were pretreated with glow discharge. n-decyl-β-D-maltoside (DM) was added to all samples at a final concentration of 0.08% (w/v) before grid preparation. Approximately 10 μM KLHL15 or CUL3-RBX1-KLHL15 mixed with 50 μM synthesized GATA2 peptide and incubated on ice for 30 minutes. Three microliters of the sample were applied to grid. The grids were subsequently blotted for 5 s (temperature at 10 ℃, and relative humidity at 100%), plunged and flash frozen into liquid ethane using a Lecia EMGP2, and stored in liquid nitrogen for data collection.

For the KLHL15-GATA2 and CUL3-RBX1-KLHL15-GATA2 complex, data collection was carried out on a Titan Krios transmission electron microscope (Thermo Fisher Scientific) operated at 300 kV at the University of Washington and the HHMI Janelia Research Campus, respectively. The automation scheme was implemented using the SerialEM software^62,63^ with beam-image shift strategy and active beam-tilt compensation at a nominal magnification of 105,000x, resulting in a physical pixel size of 0.829 Å, 0.827 Å, respectively. The zero-loss-energy images were acquired on a Gatan K3 direct detector operated in correlated double sampling mode with the slit width of post-column Gatan BioQuantum GIF energy filter set to be 20, 20, respectively. The dose rate was adjusted to 13.9, 10.1 e^-^per Å^2^ per second, respectively, and a total dose of 60 e^-^per Å2 for each image fractionated into 60 frames. The images were recorded at a defocus range of -0.8∼3.0 μm.

### Cryo-EM data processing

All datasets were processed with cryoSPARC.^64^ The beam-induced motion correction and CTF estimation was conducted with cryoSPARC using patch motion correction and patch CTF estimation. The motion-corrected micrographs were curated by defocus, CTF fitting resolution (excluding >8 Å), and ice thickness. The curated micrographs were exported and subject to template picking. After inspection of picked particles, the left particles were extracted with threefold binning and subject to 2D classification. The selected particles were subjected to ab-initio reconstruction with 4 or 3 classes. All particles then subject to heterogeneous refinement with the reconstructions generated from ab initio reconstruction as reference. Particles of good reconstruction were used as template to train model for Topaz picking.^65^

For KLHL15-GATA2, 157,009 good particles from 16,740 micrographs generated from sequential template picking, 2D classification, ab-initio reconstruction and heterogenous refinement were subjected to topaz picking. 2,426,012 particles were picked, extracted and further classified via multiple round heterogenous refinement and 2D classification. 435, 643 particles from good 2D classes were kept for heterogenous refinement. 266,513 particles were from one good reconstruction were kept and subjected to sequential non-uniform refinement with C2 symmetry implemented.^66^ These particles yield a 3.8 Å reconstruction. 2D classification was applied to further improve the local density. 236, 640 particles were kept and yield a 3.8 Å reconstruction. The following symmetry expansion and local refinement with a binary mask focused on each KLHL15 protomer domain yields a 3.5 Å and 3.6 Å reconstruction, respectively.

For CUL3-RBX1-KLHL15-GATA2, 986,150 good particles picked with topaz picking from 16,331micrographs were subjected to 2D classification and heterogenous refinement. 387,594 particles from one good class were kept for sequential particle extraction and non-uniform refinement. After refinement, 387,594 particles yielded a nominal 3.4 Å reconstruction. The densities of KLHL15 were less resolved than CUL3. To improve the density, the particles were subject to 3D classification without alignment. After 3D classification^67^, 93,113 particles were kept, which yielded a 3.6 Å reconstruction. To further polish the KLHL15 density, sequential symmetry expansion and 3D classification with binary mask focused on dimeric KLHL15. 57,689 particles were kept after 3D classification and subjected to local refinement yielded a 3.7 Å reconstruction.

### Model building and refinement

For the KLHL15-GATA2 structure, the AlphaFold predicted model was fitted into the 3.5 Å KLHL15-GATA2 EM map using UCSF Chimera^68^. The fitted model was used as the initial structure for rebuilding. For the flexible regions of KLHL15 and GATA2, the predicted model was trimmed based on the density in Coot^69^. The curated model was refined with Phenix^70^ and Rosetta^71^.

For the CUL3–RBX1–KLHL15–GATA2 complex, the KLHL15–GATA2 model and the CUL3 model from PDB 8KHP were used as starting models and docked into the cryo-EM density map using UCSF Chimera. The trimmed CUL3 model was refined against the density map using Phenix real-space refinement. The docked KLHL15–GATA2 model and the resulting composite model were subjected to rigid-body real-space refinement in Phenix. The structural figure panels were prepared with PyMol and UCSF ChimeraX.

## DATA AND CODE AVAILABILITY

Raw sequencing data generated in this study have been deposited in the NCBI Sequence Read Archive (SRA) under BioProject accession PRJNA1493822 and will be publicly available upon publication. The coordinates and cryo-EM maps were deposited in the Protein Data Bank (PDB) and the Electron Microscopy Data Bank (EMDB) with the following accession numbers: KLHL15-GATA2: 12EN, EMD-76350, EMD-76351, EMD-76381; CUL3-RBX1-KLHL15-GATA2: 12EY, EMD-76352, EMD-76353, EMD-76390.

## ACKNOWLEDGEMENT

The authors would like to thank R. Yan, Z. Zhang, N. Spellmon, and Z. Yu at the Cryo-EM Facility on the Janelia Research Campus of the Howard Hughes Medical Institute, and W. Jiang and J.D. Quispe at the Arnold and Mabel Beckman Cryo-EM Center at the University of Washington for their assistance with electron microscopy data acquisition. We also thank Y.M. Lin and N.C. Hsu of the Flow Cytometry Core, and S.Y Tung of the Genomic Core, Institute of Molecular Biology, Academia Sinica, as well as A. Silvestroni and B. Turnbull at the Department of Laboratory Medicine and Pathology Flow Cytometry Core (CC101467) at the University of Washington for technical assistance. This work was supported by the Howard Hughes Medical Institute (N.Z.), the Washington Research Foundation (B.W.) and grants AS-GCP-115-L01 from Academia Sinica and 114-2326-B-001-002 from the National Science Council of Taiwan (H.S.Y.). N.Z. is an Investigator of the Howard Hughes Medical Institute.

## AUTHOR CONTRIBUTIONS

N.Z. and H.S.Y. conceived the overall project with contributions from B.W.. H.S.Y. designed the iGPS system. C.W.Y. developed the iGPS assay. C.W.Y., H.L. and S.C. performed the iGPS screen of IDR mutations in human transcription factors. K.Y. and C.H.Y. designed the oligonucleotide library and performed statistical analyses. H.L. mapped the ZNF711 and GATA2-S447R degrons by mutagenesis. C.W.Y. performed CRISPR-based screens that identified LRRC41 and KLHL15 as the E3 ligases targeting the ZNF711 and GATA2-S447R degrons, respectively. B.W. and H.M. purified KLHL15 and characterized its interaction with GATA2-S447R. B.W. and H.S. determined the cryo-EM structures. B.W. performed mutagenesis analysis of KLHL15 mutants. B.W. identified and characterized additional proteins with FRY-like degrons created by disease mutations. B.W., N.Z., and H.S.Y. wrote the manuscript with inputs from all authors.

## DECLARATION OF INTERESTS

N.Z. is one of the scientific cofounders of and have financial interests in SEED Therapeutics and Molecular Glue Labs. N.Z. serves as a member of the scientific advisory board of Synthex, Differentiated Therapeutics, and Cold Start Therapeutics with financial interests. The authors declare no other competing interests.

## DECLARATION OF GENERATIVE AI-ASSISTED TECHNOLOGIES IN THE WRITING PROCESS

None.

